# Metabolically driven action potentials serve neuronal energy homeostasis and protect from reactive oxygen species

**DOI:** 10.1101/2022.10.16.512428

**Authors:** Chaitanya Chintaluri, Tim P. Vogels

## Abstract

So-called spontaneous neuronal activity is a central hallmark of most nervous systems. Such non-causal firing is contrary to the tenet of spikes as a means of communication, and its origin and purpose remain unclear. Here, we propose that non-input driven firing can serve as a release valve to protect neurons from the toxic conditions arising in mitochondria from lower-than-baseline energy consumption. We built a framework of models that incorporate homeostatic control of metabolic products–ATP, ADP, and reactive oxygen species, among others–by way of changes in firing. Our theory can account for key features of neuronal activity observed in many experiments in studies ranging from ion channels function all the way to resting state dynamics. We propose an integrated, crucial role for metabolic spiking that bridges the gap between metabolic homeostasis and neuronal function. Finally, we make testable predictions to validate or falsify our theory.

## 1 Introduction

Neurons fire for no apparent reason^1^, in sleeping^2–5^, resting^6^, and awake animals^2,7,8^, in acute slices^9–12^, cultures^13,14^, and organoids^15^. Such spontaneous activity is thought to be an integral part of neural function, but its origin remains unexplained. What’s more, conventional wisdom dictates that spikes relay information based on the inputs they receive. Non-causal activity seems at odds with such a model of efficient neural processing. Finally, spikes are costly, as energy is necessary to power them^16^. In this light it is unclear why neurons should fire in the absence of inputs when it would be less noisy and more energy-efficient to remain passive.

Energy consumption in neurons, broadly speaking, is the utilisation of ATP for i) restoring ion gradients after synaptic inputs, ii) general maintenance of the cellular machinery, and iii) the generation and transmission of spikes (Fig. 1a). Input (i) and output (iii) processing costs are substantial^16,17^, and ATP consumption can fluctuate widely between periods of input silence and intense activity. In order to quickly react to incoming inputs, neurons must near instantaneously match cellular ATP demands with ATP production. Such rapid ATP production in neurons is facilitated by their mitochondria (Fig. 1b), where the chemical energy from pyruvate is transformed in the tricarboxylic acid cycle (TCA, or citric acid cycle) and the electron transport chain (ETC) to establish a proton gradient across the mitochondrial inner membrane denoted by ΔΨ. When protons re-enter the mitochondrial matrix through the complex V (ATP synthase) of the ETC, ATP is milled from ADP molecules^18^ (Fig. 1c). This mitochondrial ATP is then exchanged with the cytosolic ADP to be consumed in processes such as Na-K pumps.

**Figure 1:**
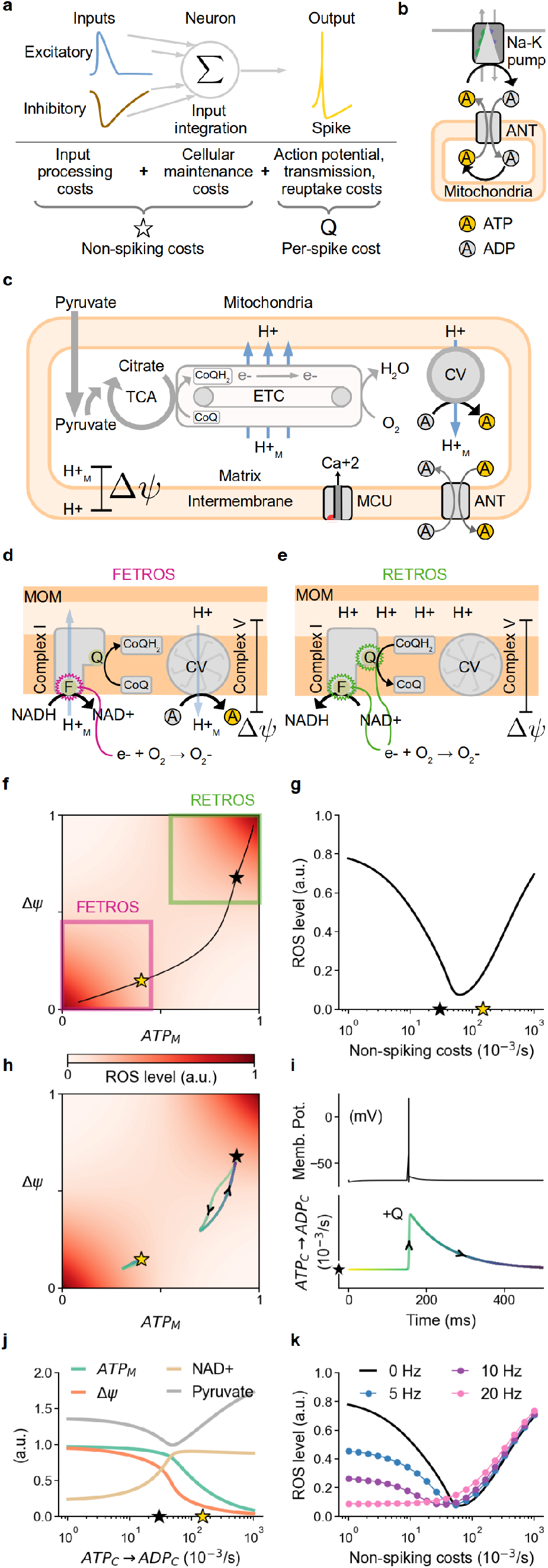
ROS homeostasis by way of metabolic spike generation. **a)** Schematic of the energy budget of a neuron, including input processing costs (left), cellular maintenance cost (middle), and action potential transmission costs (right). The sum of all input and maintenance costs is indicated by the star symbol and constitutes the non-spiking costs. The additional transient per-spike cost is indicated by Q. **b)** Metabolic energy is provided intracellularly largely through ATP, which is dephosphorylated into ADP to, e.g., drive ion pumps. This ADP is exchanged with ATP that is replenished in the mitochondria. **c)** In mitochondria, pyruvate is transformed in the tricarboxylic acid cycle (TCA, or citric acid cycle), ultimately driving protons via the Electron Tranport Chain (ETC) into the intermembrane space to generate a gradient (ΔΨ) across the inner membrane. These protons (H^+^) re-enter the matrix (H_M_^+^) through the complex V (CV) and mill ATP from ADP. **d, e)** Reactive oxygen species (ROS) production as a side-effect of proton pumping in the mitochondria: **d)** “Forward electron transport” ROS (FETROS) is routinely produced during normal function of the electron transport chain when electrons escape, e.g., at complex I during the oxidation of NADH at binding site F. **e)** “Reverse electron transport” ROS (RETROS) is produced when the electron transport chain is stalled, due to lack of ADP. Stalled electrons escape, e.g., at ubiquinone-binding site Q. **f, h)** The ‘respiratory state space’ of the extended mitochondrial model is spanned by ATP (x-axis) and ΔΨ (proton gradient, y-axis), which determine ROS production (colormap). The black and gold star mark the corresponding steady state values of ATP_*M*_ concentration and ΔΨ for two exemplary metabolic states used throughout this study. **f)** Areas of FETROS and RETROS production are marked in magenta and green squares, respectively. The black line describes the range of baseline costs. **g)** ROS levels depend on the baseline (non-spiking) costs and corresponds to the line from **f. h)** Transient trajectory away from the baseline (gold and black stars) as a consequence of a spike. **i)** Transient increase in ATP consumption over time (bottom) as a result of one spike (top). Black star in the bottom plot indicates the baseline cost. Spike-induced excursions in respiratory state space for this spike are shown in **h. j)** The concentrations of mitochondrial ATP (green), NAD+ (tan), ΔΨ (orange) and pyruvate (grey) for different steady states. **k)** Additional spiking affects ROS levels.

In all cells, not just neurons, ATP production in mitochondria routinely leads to the formation of toxic reactive oxygen species (ROS) when electrons escape from the ETC. Such rogue electrons cause free radicals which intervene in biochemical pathways and disrupt cellular processes^19^. ROS levels increase with increased number of electrons transiting the ETC in a process known as “Forward Electron Transport” ROS (FETROS) release^20^. This occurs during periods of increased ATP production that matches high ATP demand. Under these circumstances, any established ΔΨ is used to produce ATP, which in turn is rapidly utilised and turned to ADP. FETROS occurs under high ATP demands and coincides with low ΔΨ and low ATP concentrations (Fig. 1d, f, lower left magenta square).

Critically, decreased energy demands, i.e., lowered ATP consumption, can also increase ROS release^20^. This occurs when ATP is not consumed at expected rates and mitochondrial ADP becomes scarce at the complex V. Consequently, ATP production nearly stalls and the electrons lingering on the ETC escape to produce so-called “Reverse Electron Transport” ROS (RETROS)^20,21^. RETROS increases with increased ETC stalling and coincides with high ΔΨ and high ATP concentrations (Fig. 1e, f, upper right green square). Such increased ROS production in case of both increased and decreased ATP production leads to a non-monotonic, V-shaped relationship between ATP consumption and intracellular ROS levels^22^ (Fig. 1g). ROS is toxic for most cellular processes. For example in skin cells, ROS-induced stress leads to cell death. However terminally differentiated post-mitotic cells, i.e., cells that cannot be replaced or replicated such as skeletal muscle cells, are equipped with additional mechanisms to minimise ROS by scavenging the escaped electrons^23^. They may even avoid ROS formation altogether by controlling the intake of glucose – the primary 6-carbon chemical energy supply in cells – and tightly regulating its breakdown to pyruvate via glycolysis, or by storing surplus as reserves in the form of glycogen or lipid droplets.

**Central assumptions and hypothesis of our study**

#### Assumptions - (I)

In neurons, the rate limiting step of ATP production is located at its last step, i.e., complex V (ATP-synthase) of the electron transport chain in their mitochondria. **(II)** Neurons, depending on their physiological operational conditions, can face a range of metabolic demands and consequently may be exposed to non-monotonically varying ROS levels. At ideal metabolic preparedness, ROS levels are minimal and increase both when higher (FETROS) and lower (RETROS) energy demand occurs.

#### Hypothesis

Neurons sense their metabolic state and modulate their excitability to achieve metabolic homeostasis, thus minimizing ROS. Towards this goal, neurons may initiate *metabolic spikes* under RETROS conditions and limit their spiking during FETROS conditions.

Neurons are special. Despite their extensive energy demands, healthy, mature, i.e., terminally differentiated and post-mitotic neurons are not known to maintain long-term energy reserves^24–27^. Moreover, glucose intake and its breakdown via glycolysis are less well regulated in neurons^28–33^ and continuously augmented by supplements from external sources^27,34,35^ (For more details on metabolism in neurons see Supp. 1, Fig. S6, S7 and for metabolic differences between neurons, astrocytes and skeletal muscle cells see Table 3.5). These observations lead us to the central assumption of our work – in neurons, ATP production follows a lean production cycle with the rate limiting step at the last site of the electron transport chain. In other words, the rate of ATP production is controlled by (ΔΨ and) the availability of ADP at ATP synthase (complex V) of the ETC. Functionally, this is sensible because neurons need to retain a high level of metabolic readiness in order to process potentially life-threatening inputs and consequent rapid surges in ATP demands. Keeping the rate-limiting step of ATP production at the complex V can guarantee such readiness. Only a sufficiently high proton gradient and ADP would thus be required to produce ATP. Mechanistically, our assumption is congruent with neurons’ lack of long-chain carbon reserves, their reliance on leaner carbon molecules and on mitochondria for their energy requirements. The brim-full proton gradient that is beneficial to guarantee quick and reliable ATP availability would come at the cost of RETROS-poisoning during periods of synaptic quiescence when ATP production stalls^22^. It is conceivable that under RETROS conditions many ATP consuming processes such as phosphorylation, transport of cellular cargo and cellular upkeep would be enhanced, but given that up to 80% of a neuron’s ATP budget may be allocated to input processing^17^, a more comprehensive solution, in addition to ROS scavenging, seems imperative to avoid acute RETROS build-up. (For a list of ROS relieving mechanisms see Table 3.4)

Here, we propose that some experimentally observed “spontaneous” firing is in fact not spontaneous, but serves as a release valve to protect neurons from toxic reactive oxygen species that appear due to ATP-production stalling when energy consumption is low, i.e., when RETROS conditions prevail. Further, we show that such *metabolic spikes* may be behaviourally relevant and we explore the validity and the implications of our theory in 5 increasingly detailed, biologically plausible models. We begin with a mitochondrial model that links the TCA and ATP production with ROS release to show that metabolic spikes, i.e., spikes that serve metabolic homeostasis and save neurons from ROS, could provide the necessary quick increase in ATP expenditure and limit RETROS. Next, (II) we account for the frequency and the firing pattern of metabolic spikes that would be necessary to perform ROS homeostasis and explore potential mechanisms that can detect a neuron’s *metabolic state*, and drive the general changes in intrinsic firing. To showcase a single ion channel type’s response to ROS more concretely, (III) we model the reported intrinsic firing changes in dorsal fan-shaped body neurons due to *β*-subunits of Shaker channel of fruit fly (*Drosophila melanogaster*). Further, we show that metabolic spikes may not just contribute noise, but may be integral for neural function and behaviour. For this, we examine (IV) the consequence of *metabolic spikes* in a large-scale recurrent network model and compare the resulting sustained activity with previous experimental observations. Finally, we discuss the implications of our theory for a neuro-degenerative disease (V) whose hallmark symptom is altered spontaneous activity, i.e., Parkinson’s. We end our study with a set of experimentally testable predictions (VI) that would prove or falsify our theory.

## 2 Results

In neurons, several cellular processes consume ATP. In our study, we separated all ATP consuming processes into two categories based on how quickly these ATP costs can change. We consider the ATP budget for all input processing and cellular maintenance as a slow-changing, steady-state expense. We refer to this budget as the non-spiking cost or baseline cost and mark it as a star (⋆) in all figures. At baseline, the ATP-producing mitochondrial processes will match the ATP demands and arrive at steady state values of ATP concentration and ΔΨ (Fig. 1a). Secondly, we consider the ATP budget that is required to re-balance the Na-K ions, neurotransmitter release and re-uptake after an action potential as a fast transient increase from the baseline costs. We denote this budget as per-spike cost (Q) (Fig. 1a, i).

To visualise the baseline costs and the transient excursions due to a spike, we introduce the ‘respiratory state space’ (Fig. 1f, h) - a two dimensional space spanned by ATP on the x-axis and ΔΨ on the y-axis. Any given neuron will be situated at a specific location in respiratory state space according to its current baseline cost (Fig. 1f). Spikes will produce transient excursions along trajectories that depend on a neuron’s position in the state space and the per-spike cost expense Q. The neuron’s momentary position thus reflects its metabolic state.

### I. Metabolic spikes can quench mitochondrial ROS

Periods of low ATP consumption are known to lead to elevated ROS levels in other cell types but they have never been observed in neurons, despite their widely fluctuating energy consumption. We wondered if spike initiation could serve as a homeostatic mechanism that regulates ROS by restoring ATP production when energy consumption is low. To explore this idea more quantitatively, we developed a theoretical framework in which the metabolic state affects neural excitability. We started by expanding an existing mitochondria model^36^ to include ROS production and its scavenging such that ROS amplitude was a direct consequence of the concentration of mitochondrial ATP (ATP_*M*_) and the proton gradient ΔΨ (Fig. 1f-h, j, Methods 2.1-2.4).

ROS concentrations could thus be regulated by eliciting “metabolic spikes”, prompting excursions from their momentary metabolic state (black star in all figures) in the ‘respiratory state space’ (Fig. 1f, h, S1b,c). These excursions (Fig. 1h, i) from non-spiking baseline transiently increase cytosolic ATP (ATP_*C*_) consumption and thus provide mitochondrial ADP (ADP_*M*_). Depending on a neuron’s position in respiratory state space, additional spikes can decrease or increase ROS (Fig. 1h, black and gold star, respectively) independently of other variables (Fig. S1, S2). To minimise FETROS, a neuron may thus have to decrease spiking, and save ATP. To minimise RETROS, a neuron may instead have to produce spikes at various intervals and thus spend ATP (Fig. 1k). Because ROS production is non-monotonic with ATP consumption, there exists an ATP expenditure rate 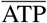 between the RETROS and FETROS conditions such that ROS levels are at a minimum (Fig. 1g).

### II. The firing patterns a neuron produces will depend on the details of its energy budget

In a minimal neuron model that tracks the metabolic costs of basic maintenance, integration, and spiking (Fig. 2a) we can estimate the firing patterns a neuron must elicit to reach a ROS minimum. In a real neuron, the metabolic state could be read out from its ROS levels directly (see below), but also from its ATP/ADP ratio, its intracellular Ca^2+^ concentration, and the cumulative footprint of its ROS scavenger response with its resulting redox potential. For the simplicity of our model, we considered only two of these variable and calculated a non-ambiguous “metabolic signal” as MS = ROS×∂ATP, where 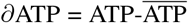 (Fig. 2b).

**Figure 2:**
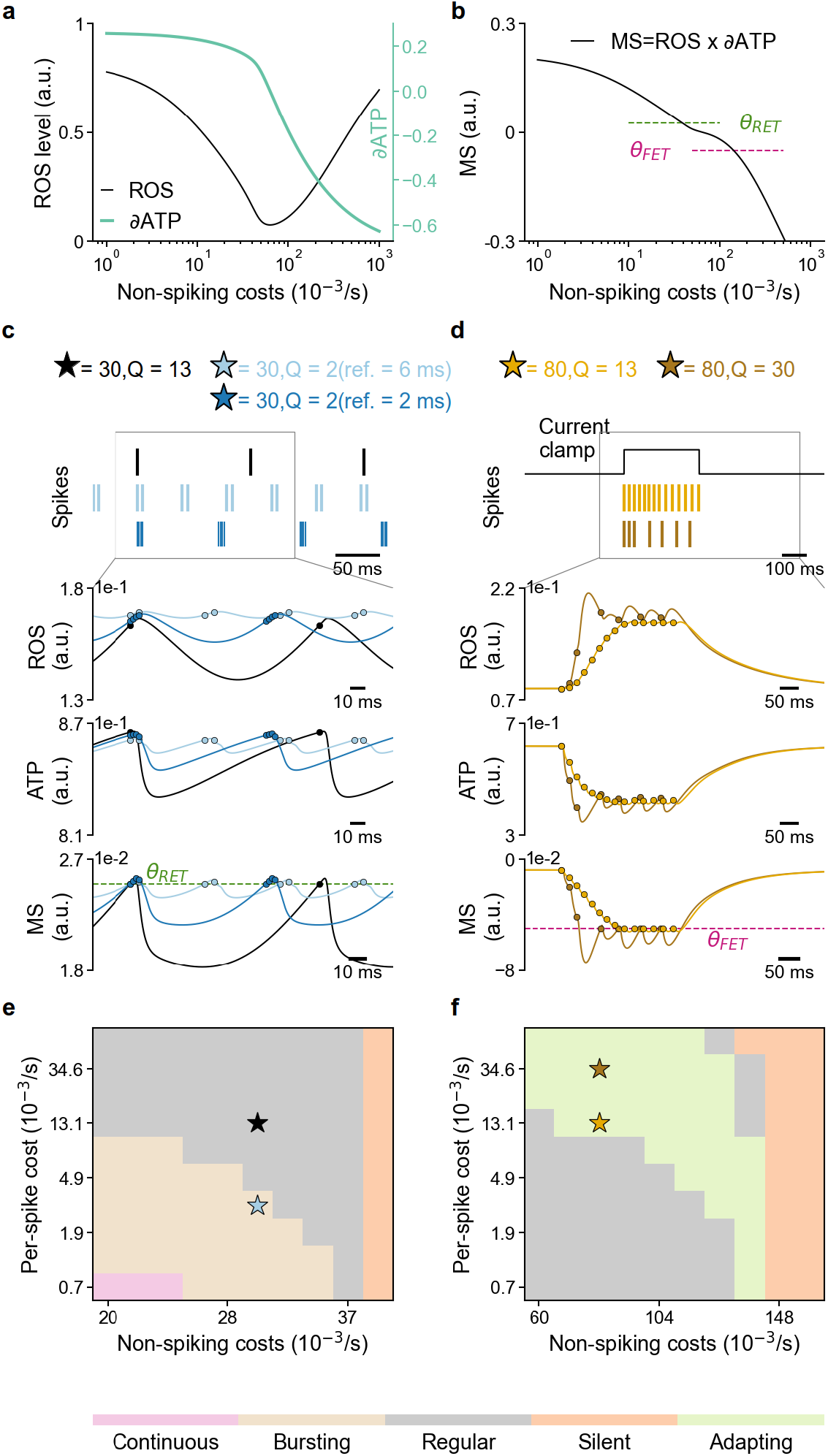
ROS homeostasis is linked to neural excitability. **a)** ROS and ∂ATP levels change with non-spike related cytosolic ATP_*C*_ consumption in our simplified metabolic accounting model. **b)** The metabolic signal (MS) as the product of ROS and ∂ATP. Two thresholds *θ*_*RET*_ (dotted green line) and *θ*_*FET*_ (dotted magenta line) control the neural response. **c)** Top panel: Raster of metabolic spikes initiated when the MS >*θ*_*RET*_ (black). When Q is low, MS remains higher than *θ*_*RET*_ after the refractory period and additional metabolic spikes are initiated (light blue). Lowering the refractory period increases in-burst frequency, and decreases inter-burst frequency (dark blue). The lower panels show the ROS, ATP and MS levels for 120 ms of the simulation (second, third and fourth row, respectively) for the three cases. Spikes are marked as ‘o’. **d)** Top panel: Raster of spikes (gold) in response to an external current source (black). When Q is high such that MS decreases and stays < *θ*_*FET*_ for longer (brown) where spiking is prohibited. The lower panels show the ROS, ATP and MS level (second, third and fourth rows). Spikes are marked as ‘o’. **e, f)** The non-spiking related baseline ATP usage (x-axis) and the per-spike-cost Q (y-axis) determine the spectrum of intrinsic firing responses ranging from silent to continuously spiking neurons in RETROS **(e)** and FETROS **(f)** conditions.

In this accounting model (Methods 2.5) the necessary spike changes to minimize ROS can be obtained by introducing two thresholds, *θ*_*RET*_ and *θ*_*FET*_. When MS is large (high ROS, high mitochondrial ATP_*M*_), such that MS> *θ*_*RET*_, the model elicits ATP-consuming, RETROS-quenching metabolic spikes (Fig. 2c). If MS decreases, such that *θ*_*RET*_ *>*MS*> θ*_*FET*_, metabolic spiking ceases. When MS decreases further (depleted mitochondrial ATP_*M*_, high ROS), such that MS*< θ*_*FET*_, even input-driven, synaptic spiking is prevented (Fig. 2c). In our model, intrinsic excitability is thus governed by MS and serves as a homeostatic mechanism to minimize ROS.

The effectiveness of a spike to modulate ROS depends on a number of variables, most prominently the transient increase in ATP consumption after each spike, i.e., per-spike-cost (Q) (Fig. 1a). When the per-spike-cost Q is sufficiently high, single spikes delivered at low rates suffice to quench RETROS. For lower Q, several spikes are necessary to decrease the MS and quench high ROS (Fig. 2c). Q also affects the FETROS response, as expensive spikes will deplete cellular ATP more and lead to prolonged periods of blocked spike initiation (Fig. 2d). The various possible combinations of Q and steady-state ATP consumption can account for many different spike different spike patterns, ranging from silent to regular firing to bursting, to continuously active (Fig. 2e, f). In mature neurons, Q would presumably be constant, but the steady state non-spiking expense (star in all figures) may change, e.g., due to changes in input activity, a neuron’s size or its history (see Fig. S3 for additional relevant variables). The firing patterns that emerge from these combinations of constraints can range from an occasional single spike to bursting or even continuous spiking and mirror many of the experimentally observed intrinsic activity patterns (Discussion).

Any energy homeostatic mechanism must be sensitive to the current metabolic state of the neuron. We propose that, excitability of the neuron must dynamically change in response to the metabolic status, and provide relief from both FETROS and RETROS. A candidate mechanisms to achieve such a swift modulation in excitability would likely involve the many ion channel types that can sense the metabolic state via metabolic signals (Fig. 3a). We can broadly categorise such metabolic state-sensing ion channels as ATP ‘savers’ and ‘spenders’ in a simplified neuron model (Methods).

**Figure 3:**
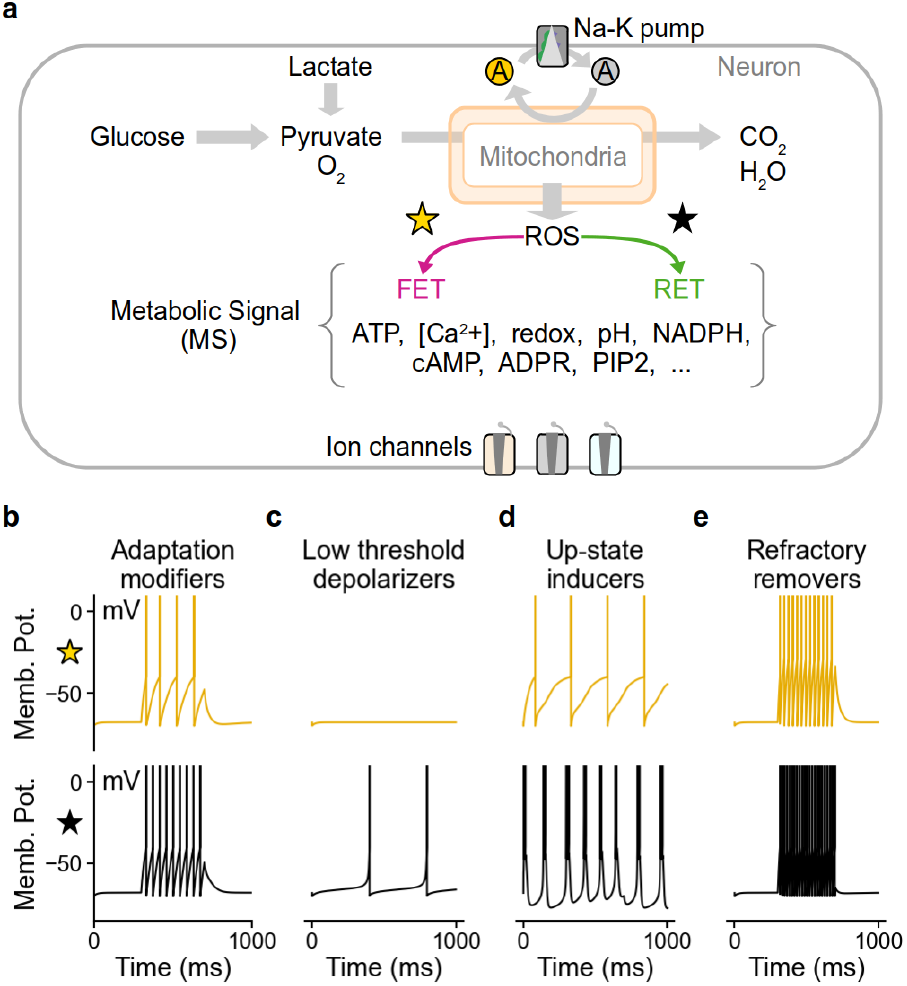
ROS homeostasis could be mediated by ion channels. **a)** ROS, its scavenging response by the cell, concentration of various substrates, its redox state, Ca^2+^ concentration, pH, etc, determine the cellular metabolic state (position of star in Fig.1j) and cause changes in ion channels or their sub-units to modulate neuronal excitability. **b-e)** Schematic of four discernible firing pattern changes mediated by ion channel modifications. **b)** Increased participation of A-type channels in spike repolarization can facilitate high frequency firing by diminishing spike adaptation. **c)** Opening of low threshold depolarising channels (e.g., persistent Na or TRPM2 channels) and the closure of hyperpolarizing channels (e.g., ATP-dependent K channels and SK channels) can cause metabolic spikes. **d)** Increased conductivity of T-type Ca^2+^ channels can promote bursting e.g., by limiting Na channel inactivation. **e)** Modification of ion channels sub-units can eliminate refractory period between spiking (e.g., seen as a consequence of resurgent Na currents).

ATP savers aim to silence neurons when FETROS conditions prevail. Their combined action delays spikes or suppress them entirely. ATP saver channels act to hyperpolarize the membrane potential such as ATP-sensitive or Ca^2+^-sensitive potassium channels (Fig. 3b top). On the other hand, ATP spenders should facilitate spiking, e.g., through channels that aid rapid re-polarisation and decrease spike-frequency adaptation (Fig. 3b bottom) or by low-threshold-activating depolarising channels (Fig. 3c). Similarly, channels that would increase bursting (Fig. 3d), or shorten the refractory period and thus effectively eliminate interspike intervals would promote ATP spending and support the firing patterns required to maintain a baseline level of ATP expenditure (Fig. 2e light purple, Fig. 3e).

Cumulative changes in many ion channels, such as changes in voltage sensitivity, activation or inactivation profiles, time constants and conductivity, and several other post-translational modifications in response to the metabolic state, will determine if a neuron decreases (Fig. 3b-e, upper panels), or increases (Fig. 3b-e, lower panels) its excitability (Discussion). Importantly, the down-regulation of ATP savers during RETOS could effectively permit ATP expenditure, and vice versa. As such, the designation of an ion channel as an ATP saver or spender would not be based on the specific ion they gate, but on the cumulative change in the firing property of the neuron. Notably, we can identify more than 15 candidate pathways (Supp. Table 3.2) that target ion channels which could counter RETROS or FETROS conditions.

### III. Linking ion channel function with metabolic state and ROS

One example of using spikes to sustain the ETC has been observed in dorsal fan-shaped body (dfb) neurons of *Drosophila melanogaster*^37^(Fig.4a). In these sleep-switching dfb neurons, the A-type “Shaker” potassium channels’ *β*-subunits called “Hyperkinetic” are modulated by a key molecule involved in ROS scavenging called nicotinamide adenine dinucleotide phosphate (NADPH) (Fig.4b). When ROS is high, NADPH is converted into NADP+, neutralising ROS, but also eliminating the channels’ inactivation mechanism. The fast-activating, fast-inactivating Shaker-Hyperkinetic channel thus changes to slow-inactivating during RETROS states (Fig.4c). This loss of the inactivation gate of a K^+^ channel is thought to be the cause for a behaviourally relevant increase in firing (Fig.4f, top).

**Figure 4:**
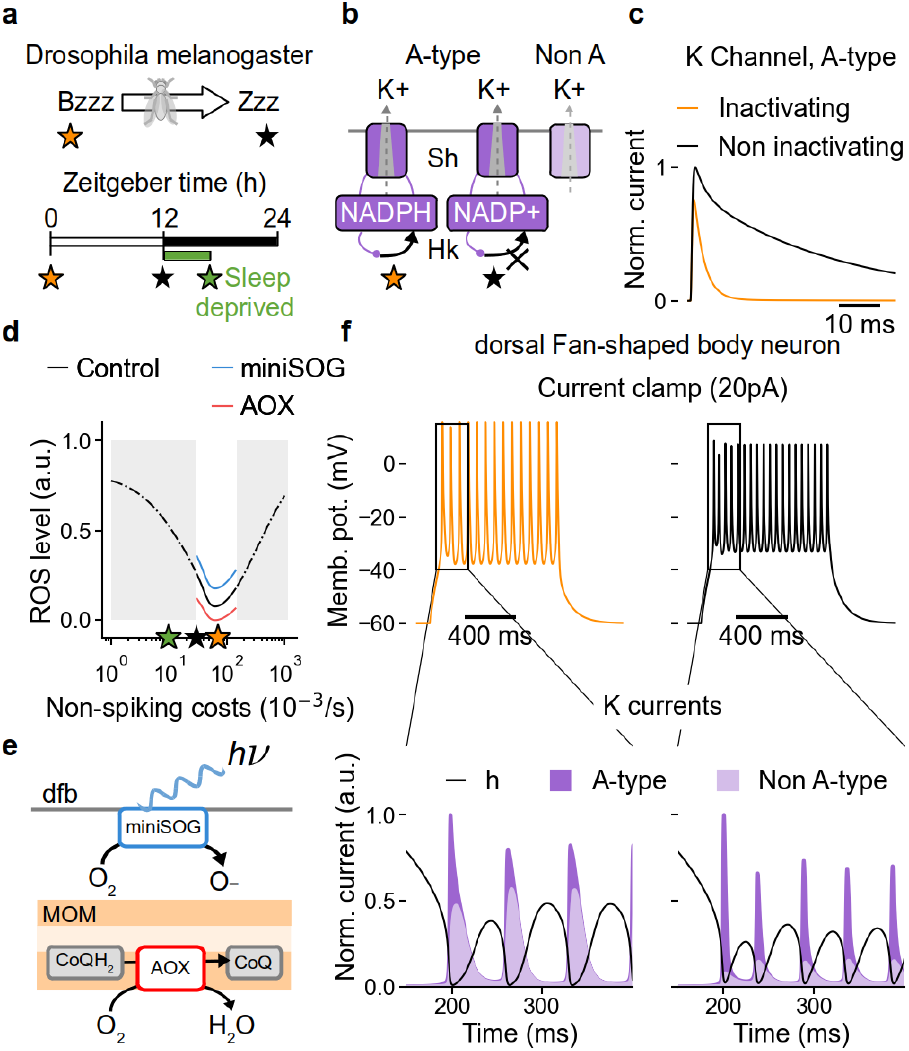
Functional consequences of ROS homeostasis. **a)** Changes in intrinsic firing of the dorsal fan-shaped body neuron in fruit flies may be related to changes from high (orange star) to low metabolic state (black star) and switch the behavioural state from awake to sleep^37^. **b)** The origin of firing changes is thought to lie in the altered inactivation mechanism of NADPH-receptive Hyperkinetic *β*-subunits (Hk) of Shaker (Sh) channels. Changing their profile from inactivating (orange star) to non-inactivating (black star) affects the ratio of A (dark purple) to non-A (light purple) type potassium currents. Voltage clamp response of inactivating (orange) and non-inactivating (black) A-type currents for a 40mV step. **d)** Changes in ROS production (black line) as a function of non spiking costs. The neurons’ presumed physiological range is indicated in white background. Sleep deprived state with its hallmark of high firing and increased ROS levels lies outside of the physiological range (green star)^37^. ROS production can be altered by way of miniSOG (blue) and AOX (red). **e)** miniSOG increases ROS optogenetically (top); overexpressing AOX oxidizes coenzyme-Q and decreases ROS levels (bottom). **f** top) Spiking response to a 20pA current injection in neurons with inactivating (orange) and non-inactivating (black) K current; Bottom) zoomed-in visualisation of the (inactivating (left), or non-inactivating (right)) A-type (dark purple) and non-A-type (light purple) K current contributions, co-plotted with the sodium current’s inactivation variable h (black).

We can support this assumption and reproduce ROS-dependent firing changes in a standard Hodgkin-Huxley type model with an additional A-type K^+^ channel, resembling Shaker-Hyperkinetic channel, that alters its inactivation properties in response to ROS. In our model (Methods 2.6), slowed inactivation of the A-type K^+^ channel leads to increased participation of its currents in membrane re-polarisation, out-competing slower-activating, non-A-type currents and thus allowing for more rapid re-polarisation (Fig.4f, bottom) and consequently decreased inter-spike intervals (and thus an increase in firing) as consistent with experimental observations.

In the experiment this can be directly linked to ROS because, when the fruit fly is sleep deprived (Fig. 4d green star), the dfb neurons are pushed beyond their physiological operational preparedness (Fig.4d grey background). In our framework this can be interpreted as a situation in which metabolic homeostasis through spiking becomes insufficient and the additional RETROS burden is revealed. In the experiment, a higher ROS burden can also be delivered optogenetically through mini Singlet Oxygen Generators (miniSOG) and can this be simulated congruently by an upward shift in the relationship between baseline ATP consumption and ROS concentrations (Fig. 4d, e blue). Conversely, a ROS decrease can be achieved experimentally by Alternative Oxidase (AOX), providing an additional exit for electrons from the ETC (Fig. 4d, e red). In the model this can be simulated accordingly by a downward shift in the relationship between baseline ATP consumption and ROS concentrations (Fig. 4d). We wondered if such consistency between model and experiment could also be observed in other neural systems.

### IV. Metabolic spikes in a recurrent neural network model

To explore the effects of spike-induced energy homeostasis in neural circuits, we built a network model^38^ of 8,000 excitatory and 2,000 inhibitory conductance-based leaky integrate-and-fire neurons. We used previously published neuron and network parameters, such that each neuron received recurrent inputs from a random 2% of the network as well as from external Poisson inputs. Additionally, each neuron is subject to a metabolic current *I*_*M*_ that was controlled by a metabolic state variable MS (Fig.5a, Methods 2.7) tracking the metabolic costs of synaptic inputs and action potentials. MS controlled the metabolic current *I*_*M*_ such that high MS produced depolarising membrane currents, and low MS lead to hyperpolarising currents (Fig.5b). When the network receives external stimuli, it initially fired at high rates and thus utilised large amounts of ATP, consequently operating in the FETROS regime with very low MS values. As a result, the MS-induced metabolic current becomes hyperpolarizing, contributing to a stable balance of excitation and inhibition and lowering firing rates to prevent seizure-like states (Fig.5c-f, left). Such dampened activity shifts the metabolic load near the ROS minimum. When external inputs are turned off, recurrent synaptic activity alone cannot self-sustain indefinitely, and network activity collapses (Fig.5c, d, end of yellow bar). Due to the lack of synaptic input, and subsequent decrease of ATP consumption, the MS controlled current becomes depolarising, eventually causing metabolic spikes (Fig.5c-f, middle). These first few spikes in the network rapidly activate their similarly depolarised downstream neighbours, and eventually activate the entire network, preventing RETROS build-up via synaptic currents and spiking (Fig.5c-f, right). Depending on various variables such as per-spike cost, as well as the structure of the network, activity may briefly cease again (Fig.5c-f, right), before experiencing the next RETROS induced *avalanche*^12^.

Such avalanches are indistinguishable from externally driven activity (Fig.5g-i), except for their internal metabolic status (Fig.5j). The duration of the activity exhibits a power-law distribution, similarly to widely observed *spontaneous* activity in experimental set-ups^12,39^ (Fig.5k-m). Despite the absence of external inputs, the network responded with stable and robust activity for a wide range of synaptic parameters (Supp. Fig.S4). The range of synaptic strength values for which the network displays asynchronous and irregular activity is substantially bigger than reported in previous studies^38^ (Fig.5c-f, left, Supp. Fig.S4 a-c)). In fact, as long as the metabolic currents were not maxed out (and overwhelmed by synaptic currents, Supp. Fig.S4 g), MS allowed the network to self-tune its population rate (Supp. Fig.S4 a) and asynchronicity (Supp. Fig.S4 c, e) to remain persistently and indefinitely active even for parameters settings that would otherwise lead to severe instabilities (Supp. Fig.S4 b,d,f,j). Beyond the implication in avalanche-like activity in neuronal networks (Fig. 5g-m), metabolic spiking can thus provide sustained, network-wide, asynchronous irregular activity that is thought to be the basis of robust neural circuits^40^.

**Figure 5:**
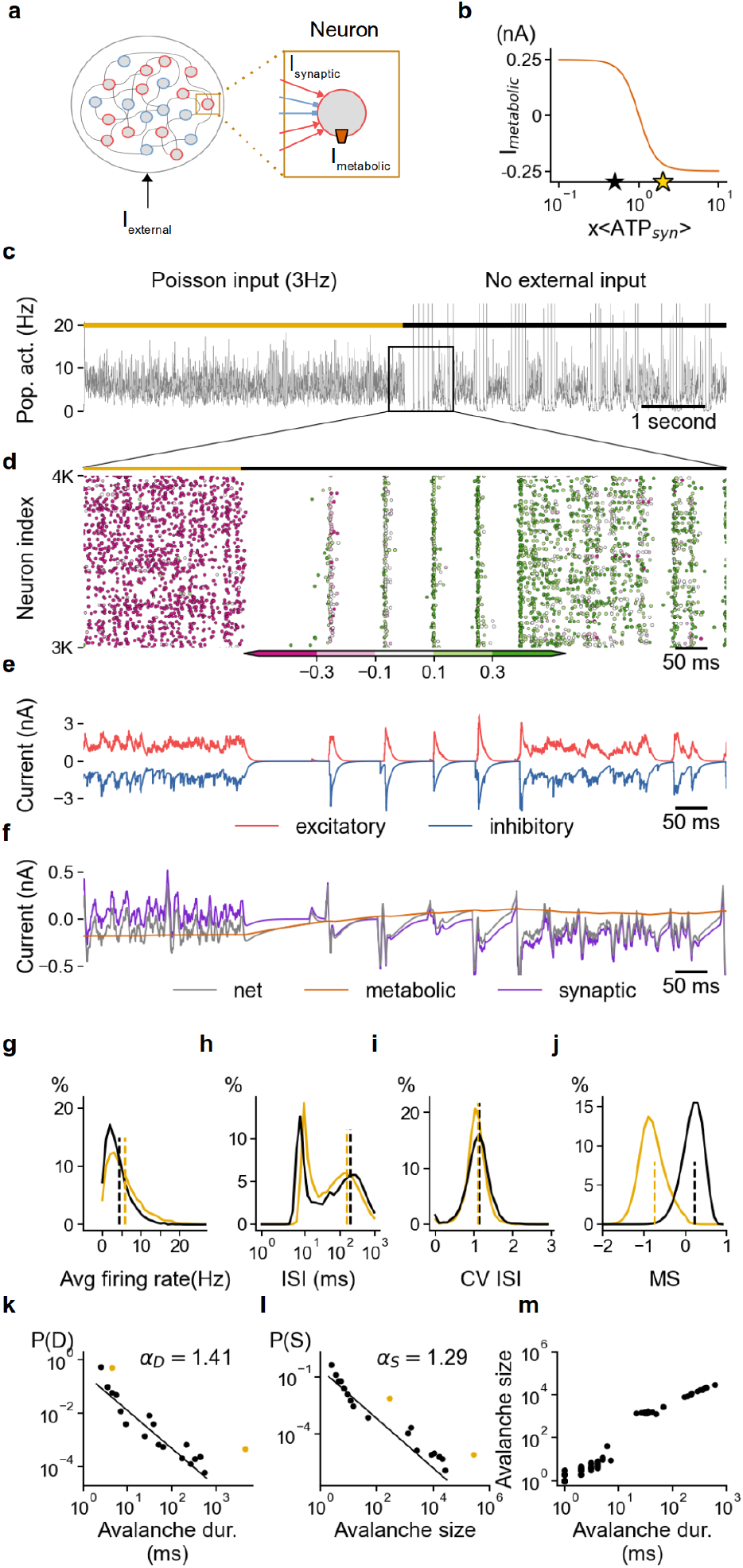
Functional consequences of metabolic spikes in a recurrent network. **a)** Schematic of a recurrent network model with excitatory (red) and inhibitory (blue) neurons that are equipped with a metabolic spiking mechanism (orange box in the single-cell inset, middle). **b)** I_metabolic_ contributes a homeostatic current according to a neuron’s energy consumption due to synaptic currents I_synaptic_. **c)** Network population rate over 10s in 1ms bins, with (gold bar) and without (black bar) external Poisson input. **d)** Spike raster inset of 1000 randomly chosen neurons during the transition from externally driven spiking to internally generated activity. The colour of each spike reflects the metabolic state of its neuron, ranging from ATP-depleted (magenta) to ATP-rich (green). Excitatory (red) and inhibitory (blue) synaptic currents of an example neuron during the transition. **f)** 5ms moving average total synaptic current (purple), metabolic current (orange) and net sum of all currents (grey) during the transition for the neuron in e. **g - j)** Distributions of neuronal firing rates (**g**), their interspike intervals (ISI, **h**) coefficient of variation (CV) of ISIs (**i**), and metabolic signal at time of spiking (**j**) before (gold) and after (black) the transition. The dashed lines show the mean of each distribution. Distribution of duration (**k**) and size (**l**) of continuous activity (separated by at least 1ms of silence) in the network before (gold) and after (black) the transition. Poisson-driven activity remains ‘on’ forever. The duration of metabolic ‘avalanches’ follows a power law distribution (black line). **m**) The avalanche size and its corresponding duration are also shown.

### V. Loss of metabolic spikes in Parkinson’s disease

Next we argue that if metabolic spikes are incorporated into the basic framework of neural function and provide a guaranteed baseline of activity, they may offer a new perspective on neurodegenerative disease, and specifically Parkinson’s. In this disease, the dopaminergic neurons in the substantia nigra pars compacta (SNcDA) cease to fire and eventually die, with catastrophic consequences: the silence of these normally tonically active neurons with long axons and dense, cotton-ball like arborizations (Fig. 6a, Fig. S5) leads to depleted dopamine levels in the striatum^5,41,42^, consequently disrupting the gating pathways of the basal ganglia^43^.

**Figure 6:**
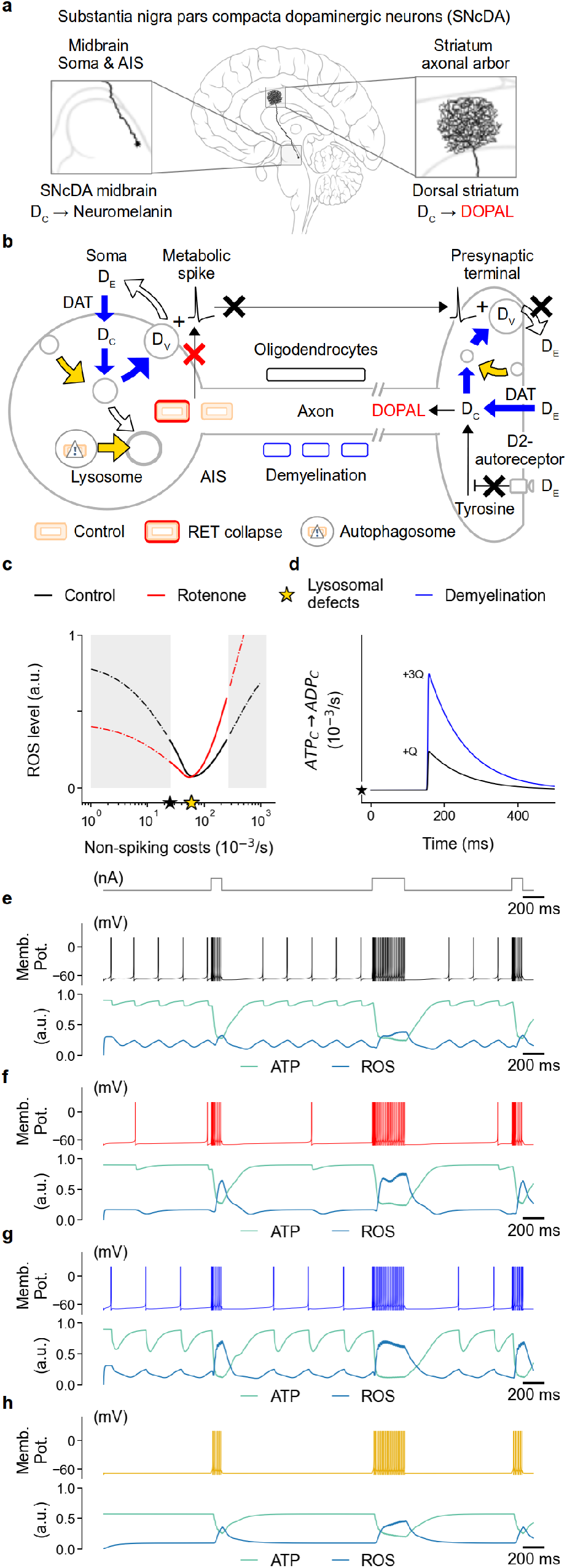
Loss of metabolic spiking may cause the symptoms of Parkinson’s Disease. **a)** Degeneration of substantia nigra dopaminergic (SNcDA) neurons as a result of decreased “tonic” firing and concurrently diminished dopamine release in the striatum disrupts the direct and indirect pathways of the basal ganglia. SNcDA neurons have long, cotton-ball like axonal arbours in the striatum. **b)** Decreased tonic spiking of SNcDA neurons could arise from 3 types of deficiencies in metabolic spike generation: First, collapse of RETROS production, or failure of ROS detection in the spike initiation zones (shown in red). Second, increases in per-spike-cost Q may arise from loss of myelin (blue boxes), poor synaptic vesicle loading and release and re-uptake of dopamine (blue arrows). Third, increased overall baseline costs, e.g., due to inefficient endosome formation or poor lysosomal function, results in a general shift of non-spiking baseline costs towards FETROS (denoted with gold arrows). In all the above cases, decrease in metabolic spiking and hence diminished extracellular dopamine (black crosses), may lead to a compensatory up-regulated dopamine production, by way of the D2-autoreceptor, that may contribute to deleterious DOPAL in the cytosol. **c)** The neurons’ presumed physiological range is indicated by the white background. Changes in the ROS levels under control/healthy (black line) and due to RETROS collapse e.g., due to chronic rotenone exposure (red line). The stars indicate the baseline ATP expenditure for the case of control (black star) and poor lysosomal function (gold star). **d)** Assumed per-spike-cost Q for a SNcDA neuron is shown for the control (black line) and the case with increased per-spike-costs, e.g., due to demyelination (blue line). **e-h)** Membrane potential (top) and corresponding ATP and ROS levels (bottom) of a metabolism linked SNcDA neuron model in response to a current injection (0.3 nA) (top-most panel of e). **e)** Control/healthy neuron. **f)** Simulated collapse of RETROS levels, e.g., by chronic rotenone exposure (panel c). **g)** Increased per-spike-cost Q (panel d), e.g., by loss of myelin. **h)** Increased baseline expenditures (panel c gold star), e.g., by inefficient lysosomal function.

We wondered if the cause for the lack of spiking of SNcDA neurons could be of metabolic origin. We thus built a simplified SNcDA neuron model that expanded on the previously used leaky integrate-and-fire neuron. The model neuron tracked metabolic expense and ROS production, and featured an additional ROS sensing depolarizing channel, and an ATP sensing hyperpolarizing channel (Fig. 6e). A depolarising current step drives the neuron to burst (Fig. 6e, upper two panels). When input ceases, the resulting (RET)ROS increase (Fig. 6e, lower panel) will trigger metabolic currents which in turn elevate the neuron’s membrane potential and eventually lead to (tonic) firing, in agreement with the results in previous sections (Fig. 6c-e black lines). We then studied how to perturb the model neuron’s behaviour and found three categories of deficits in homeostatic regulation which could result in diminished dopamine release:

1. RETROS collapse: We simulated a collapse of RETROS production by altering the relationship between metabolism and ROS production (Fig. 6c red line) such that less RETROS was produced for the same ATP and ΔΨ, directly leading to lowered tonic spiking in our model (Fig. 6f).
2. Increase in per-spike-cost: When we simulated an increased per-spike-cost by increasing the parameter Q in our model (Fig. 6d blue), the overall ATP consumption after a spike rose, making additional *metabolic spikes* unnecessary for longer periods of time (Fig. 6g).
3. Increase in non-spiking costs: Thirdly, we simulated an increase in the baseline non-spiking expenditure in our model (Fig. 6c gold star). Subsequently, as the baseline available ATP was lower, RETROS was diminished and no *metabolic* spikes were initiated at all in our simulation (Fig. 6h).

Importantly, the immediate, direct consequence of any of these deficits was a decrease in metabolic spiking in our model neuron that would–in the biological system–lead to a much decreased dopamine tone at the postsynaptic terminals in striatum (Fig. 6b black arrows), with potentially grave downstream consequences (Discussion).

### VI. Testable predictions

We have argued that neurons experience ROS and in particular RETROS as a result of a high proton gradient in the absence of mitochondrial ADP. Spiking can serve as a means to resupply ADP to a stalled ETC and thus maintain minimal RETROS levels. Our theory is based on a considerable body of indirect evidence (Supp. Table 3.1), but because ROS is detrimental to a cell’s health, we can make a number of experimental predictions to test our theory directly. For example, Purkinje neurons are known to exhibit so called ‘simple spikes’ spontaneously (Fig. 7a, b), in the light of our framework potentially as a homeostatic response to minimize RETROS. By forcefully silencing Purkinje cells with a DC hyperpolarizing current (Fig. 7b) we would suppress such ongoing spikes. Following our theory, such a perturbation should result in increased ATP levels (Fig. 7c), increased ROS (RETROS) and accelerated cell death, confirming the survival-critical nature of these spikes.

**Figure 7:**
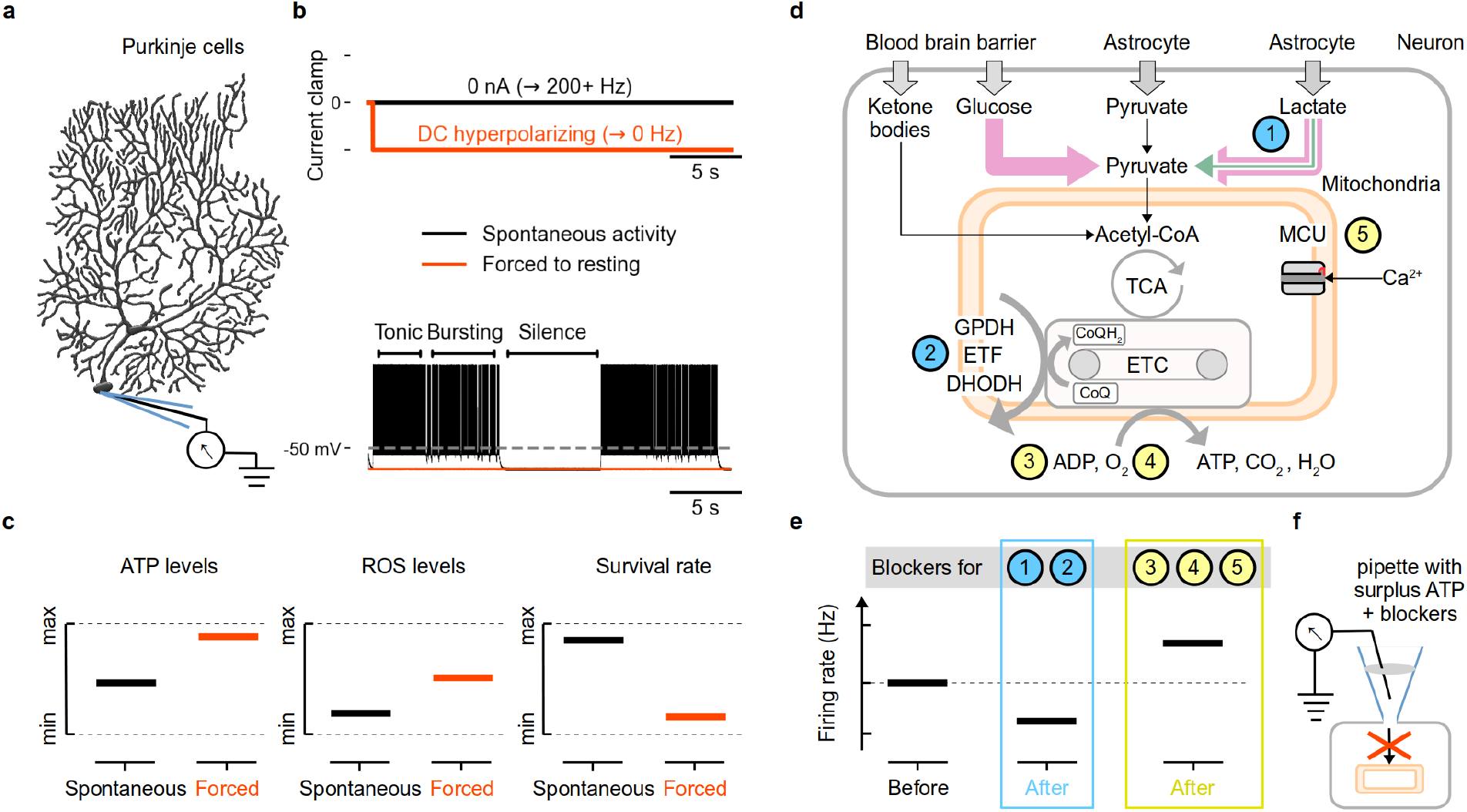
Predictions to test the metabolic excitability hypothesis. **a)** Metabolic spikes may play a critical role in the cell survival especially in neurons naturally exhibiting triphasic activity such as Purkinje cells. We predict that **(b)** forcing such spontaneously active cells to resting by DC hyperpolarizing currents would lead to **(c)** increased intracellular ATP levels and a subsequent increase (RET) ROS levels. This perturbation in the long term would decrease the survival rate of neurons. We also predict that the metabolic spikes are initiated due to ATP production stalling in neuronal mitochondria and to test this we propose experiments which decouple mitochondria from the neuron at select junctions **(d-e). d)** Schematic of the metabolic relationships in neurons, with numbered locii of possible intervention. Neurons (grey box) rely mostly on glucose under FETROS (left pink arrow) and lactate during RETROS and FETROS (right green+pink arrow) for their pruvate supply. Pyruvate is processed in the neuronal mitochondria (peach box) in their tricarboxylic acid cycle (TCA) and subsequently in the electron transport chain (ETC) to produce ATP and H_2_O by consuming ADP and O_2_. Other metabolites such as ketone bodies from blood directly enter TCA via Acetyl-CoA and circumvent pyruvate. While maintaining ample ATP in the pipette, **e)** blocking the numbered pathways (1,2) in neurons that can limit RETROS conditions may lead to a decrease in the spontaneous firing rates (blue). Conversely, blocking pathways (3-5) that would simultaneously increase the ETC stalling and ΔΨ can cause membrane potential depolarization or an increase in the firing rates (yellow). See text for more specific details. **f)** These interventions must be conducted with ample ATP supply to promote RETROS during testing.

To validate that some “spontaneous” spikes can be of metabolic origin, we propose to decouple metabolism and its hypothesised neural response during periods when the neuron is compensating for RETROS. For instance, under ATP-surplus cytosolic conditions, interventions that decrease the number of electrons stalled on ETC (or more precisely interventions that decrease the pool of electron carrying coenzyme-Q molecules (CoQH_2_) on the ETC) (Fig. 7d, 1-2), such as

1. blocking lactate dehydrogenase (LDH1/LDHB) that would decrease pyruvate production and would slow the tricarboxylic acid cycle (TCA), or
2. blocking pathways that are independent of TCA, that also transfer electrons into the ETC, such as mitochondrial glycerol-3-phosphate dehydrogenase (GPDH), electron-transferring flavoprotein complex (ETF) and dihydroorotate dehydrogenase (DHODH), should result in decreased metabolic firing rates and generally lowered neuronal excitability. According to our framework, the above perturbations (1-2) should result in a decreased RETROS production, and thus decreased numbers of metabolic spikes for the same homeostatic effect. We can also make a number of predictions for perturbations downstream from the ETC that increase the number of stalled electrons (Fig. 7d, 3-5). More precisely increasing the pool of coenzyme-Q molecules carrying these electrons on the ETC (CoQH_2_), we expect under ATP-surplus cytosolic conditions that:
3. blocking the exchange of cytosolic ADP with mitochondrial ATP at the adenine nucleotide translocator (ANT),
4. limiting O_2_ - which is the usual safe exit of the electrons on the ETC (complex IV) - while maintaining the cytosolic Na^+^ concentrations, or
5. limiting RETROS relief per-spike by blocking the mitochondrial Ca^2+^ uniporter (MCU mode 2, Discussion),

should lead to further stalling of the ETC and increased ΔΨ across the mitochondrial intermembrane space, and therefore increase RETROS. Under these perturbations, neurons should thus modulate their excitability to increase the metabolic firing.

## 3 Discussion

In cardiac mitochondria it has been shown that ROS formation accelerates during periods of increased *and* decreased energy demands^22^. Here, we postulated that neurons can manipulate their excitability to minimise the intracellular concentration of ROS. This means that when energetic demands are high, neurons may decrease spiking to avoid FETROS. Conversely, when energetic requirements are too low, metabolic spikes are initiated to produce ADP and relieve RETROS accumulation.

### I. ROS prevention through metabolic spikes

Unlike other RETROS relieving mechanisms such as ROS scavenging^23^, increased lysosomal activity or creatine phosphorylation, (See also Supp. Table 3.4), metabolic spikes can provide sustained and rapid RETROS relief that may be integrated into the existing set of neuronal heterosynaptic homeostatic mechanisms^44,45^, providing control at fast timescales^46^. We have focused on ROS and ATP as the most pertinent molecules of the neuronal metabolism, but there are likely multiple other contributors that can modulate neuronal spiking, as well as neuronal metabolic activity.

Most notably we should mention neuronal Ca^2+^, which can change dramatically due to spiking–regardless of its origin, metabolic or otherwise. In RETROS conditions, Ca^2+^ influx to mitochondria is rapidly buffered, but the additional divalent cations crossing the mitochondrial membrane would decrease ΔΨ and increase the beneficial effect of metabolic spikes. We have incorporated Ca^2+^ influx as a direct effect on ΔΨ in the relevant models (Figs. 1, 2, 6, Methods, and also Fig. S1l-n). Ca^2+^ influx to mitochondria may be mediated by the calcium uniporter (MCU in mode 2)^47–49^, which is why we suggested MCU as a possible site of intervention (Fig. 7d-5, e-5). On the other hand, the increased synaptic inputs and output spikes in the FETROS regime should lead to greatly increased cytosolic Ca^2+^ levels. Influx to the mitochondria would be slow but Ca^2+^ would remain unbound due to saturated mitochondrial buffers (MCU mode 1). Such free Ca^2+^ could serve as a signal for higher cellular energetic requirements to the mitochondria by enhancing calcium-sensitive mitochondrial matrix enzymes and increase ATP production^47,48,50^ (and thus temporarily worsen FETROS). Conversely, cytosolic Ca^2+^ would down-regulate neural excitability through Ca^2+^ dependent K^+^ channels and thus decrease spiking and hence limit FETROS. Calcium (and other variables^45^) could thus be considered part of the metabolic signals that orchestrate the responses to keep neurons safe.

### II. Ion channels as metabolic sensors and regulators

The physiological operational regime of neurons between RETROS and FETROS conditions would necessitate that they manipulate their excitability in either direction, presumably by way of several different ion channel types. Metabolic signals may not be limited to ATP, ROS and Ca^2+^ but could include signalling molecules in metabotropic cascades (Fig. 3a, Supp. Table 3.2). For example, the absence of G_*q*_PCR activators could be indicative of lowered input activity and may lead to surplus membrane-bound PIP_2_, modifying ion channels and thus neuronal excitability^51,52^. Similarly, cyclic-AMP (cAMP) concentration, which is enhanced or diminished by G_*s*_*α* and G_*i*_*α* subunits, respectively, can directly affect cAMP-sensitive ion channels such as hyperpolarization-activated cyclic nucleotide–gated channels (HCN)^53,54^.

Metabolic signals may also originate from the pentose phosphate pathway or DNA repair pathways^55^ that are favoured under RETROS conditions, when excess ATP inhibits glycolysis. Here, a potential candidate metabolic signal is adenosine diphosphate ribose (ADPR) which triggers ribosylation and thus activation of ion channels. Finally, transient extracellular acidification due to more (or fewer) ambient neurotransmitters (glutamate and GABA being abundant and acidic) could be indicative of neural metabolic load and may be detected by extracellular domains of ion channels (such as sodium) changing their pore opening probability.

A special note on the ubiquitous, low-threshold T-type Ca^2+^ channels may be warranted. Under RETROS conditions, when ROS scavenger pools operate near maximum capacity, the cytosol becomes reduced, thus increasing T-type Ca^2+^ channel conductivity^56,57^. Increased T-type conductivity in turn facilitates transient UP states that promote bursting and bursting-like firing patterns in accordance with our hypothesis. Importantly, the ion channels that allow for counter measures against ROS do not have to be de-novo expressions. Instead they are likely post-translational modifications (e.g., phosphorylation, oxidation, S-glutathionylation, ADP-ribosylation, interactions with pyridine nucleotides, phospholipid docking and so forth) that provide a response to changes in metabolic demands.

### III. Multi-channel interactions in an experimental study of RETROS

In the dfb neurons of *D. melanogaster*, ROS levels increase slowly over the duration of the day presumably due to decreasing non-spiking costs nearing sleep (Fig. 4a, b, d orange to black star). The specific reason for this decrease is as yet unknown and may involve decreased synaptic input to dfb neurons over the day. Adhering to the experimental findings^37^, our model only includes the change to the inactivation of the Shaker-Hyperkinetic channels which modulates A-type currents’ domination over non A-type currents in spike re-polarization. Consequently, the spike frequency adaptation in these neurons is relaxed under RETROS, i.e., the interspike intervals are shortened to accommodate more spikes. Such an interplay between two ion channels, with the only ROS-mediated change involving the inactivation profile of *β*-subunits, highlights the difficulty of specifying the exact mechanisms involved in metabolic spike generation in every neuron type.

Importantly, the time scale of the metabolic change in these dfb neurons is of the order of several hours. To facilitate a homeostatic response, the specific combination of ion channel and its metabolic signal must operate at speeds compatible with the speed at which non-spiking costs change in the neuron. Depending on a neurons’ placement in a circuit, and its function, we anticipate that a different combination of ion channel changes could achieve similar firing rate changes to serve metabolic homeostasis.

### IV. Sustained and finely governed network dynamics through energy homeostasis

The activity we simulated is similar to experimentally reported data seen in neural avalanches^12,39^ and in *in vivo* recordings^3,58^. At the level of populations of neurons, metabolic homeostasis guarantees baseline firing, self-governed network activity, and ensures an essential level of *input vigilance*. From first impressions, metabolic spikes may seem counterproductive, contributing noise to neural processing. However, such superfluous activity could easily be incorporated into coding schemes, serving to increase the dynamic range and sensitivity of population-based coding by providing non-zero baseline activity (Fig. 5c, d black). In previous studies, persistent activity in neural network models could only be guaranteed by way of additional external inputs. The metabolically monitored membrane currents *I*_*M*_ make such external inputs unnecessary, as the network becomes self-tuning. Further, spurious spiking of any origin, metabolic or otherwise spontaneous, can easily be cancelled out in excitation / inhibition balanced networks such that coding is not affected, whereas silent networks would be massively perturbed by initialisation and synchronisation artefacts at the onset of a given stimulus^59^. The fine control of persistent activity may promote metabolic spontaneous spiking to also serve crucial integrated functions beyond that of a safety valve preventing ROS toxicity, and it is easy to imagine tasks such as cortical waves^60^ or simple background computations rely on consistent baseline activity.

Our hypothesis of metabolic spikes as a means of metabolic homeostasis does not imply that an observed spike is exclusively synaptic or metabolic. Instead, we consider spikes to be agnostic, answering both synaptic and metabolic demands. An auditory neuron spiking at hundreds of Hz^61^ without input could be considered in metabolic spiking mode. Upon presentation of relevant input, firing transiently increases, translating synaptic input into rate code. Likewise, in Purkinje cells, neurons exhibit simple and complex spikes^62^, which maybe considered as metabolic and synaptic respectively. In addition to input integration and metabolic homeostasis, spikes may serve network functions such as a neuromodulatory state. VTA dopaminergic neurons fire at a baseline (metabolic) firing rate that either transiently increases or decreases coding for reward prediction errors^8^.

### V. Parkinson’s disease and dopamine dysregulation in SNcDA

If metabolic spikes serve a functionally integrated role, any failure to detect metabolic state and produce spiking could have system-wide, catastrophic consequences. Such a failure may be exemplified in neurodegenerative diseases such as in Parkinson’s (PD). In PD we can identify the origin of its signature symptom, i.e., the absence of dopamine in striatum^41^, as a failure to detect and respond to changes in metabolic state, and subsequent loss of metabolic spiking (Fig. S5 and Supp. Table 3.3). We can link three different failure modes for metabolic (tonic) firing in our model to well-known aspects of PD:

#### 1) RETROS collapse

In biological neurons RETROS could be diminished, e.g., via blockage of the complex I ubiquinone-binding site, which would similarly lead to decreased RETROS production (Fig. 6b red box). In line with this argument, chronic rotenone competitively binds near the complex I ubiquinone-binding site, and has been shown to reduce RETROS and cause Parkinson’s-like symptoms^20,63–65^ (Fig. 6c red, Fig. S5a). Similarly, age^66,67^ and sex^68–71^ related inefficiencies of loading electrons onto the ETC could lead to inefficient ETC and TCA, subsequently to lowered ETC-stalling, and thus RETROS production (Fig. S5b). Additionally, any abnormalities in tagging, trafficking, and degradation of mitochondria marked for repair or mitophagy^72,73^ could cause resource competition with their healthy counterparts. Such faulty mitochondria would consume resources without replenishing ΔΨ and thus do not contribute to RETROS, leading to diminished metabolic spiking (Fig. S5c). Of course, metabolic spiking could also be disrupted by ion channel mutations or other sensor failures directly^74^ (Fig. 6b red cross).

#### 2) Increase in per-spike-cost

In SNcDA neurons, an increase in per-spike-cost could happen due to, e.g., de-myelination of axons (Fig. 6b blue boxes), such that affected neurons would use more ATP per spike (Fig. 6d blue) to re-balance the Na^+^ and K^+^ ions and thus decrease RETROS more effectively, and allow them to fire less often for the same homeostatic effect. This effect may be particularly relevant in SNcDA neurons due to somatodendritic dopamine release where inefficient dopamine transmission after each spike could lead to increases in per-spike-costs near the axon initial segment (Fig. 6b blue arrows). For instance, inefficient and ineffective neurotransmitter loading into the vesicles (Fig. S5e) and spike-mediated neurotransmitter release (Fig. S5f) and re-uptake could also lead to increases in per-spike costs.

#### 3) Increase in non-spiking costs

An increase of the baseline ATP expenditure could happen through faults in lysosomal degradation and recycling of cellular components (Fig. 6b gold arrows). Defects in the somatic lysosomes which are responsible for breakdown of worn-out cellular macromolecules to nutrients could cause a dearth of available metaboloids for cellular processes. This would increase the non-spiking ATP expenditure of the neuron due to the *in situ* production of these metaboloids (Fig. 6c gold star). These faults may already arise at the ATP-expensive clathrin-mediated endosome formation pathways^75,76^ (also relevant in synaptic vesicle formation, Fig. S5g). Further, faults can arise in the transport of late endosomes and autophagopore to the lysosome, in lysosomal acidification, or faults in the co-chaperone mediated autophagy (CMA) and the recycling action of the lysosome itself e.g., by glucocerebrosidase (GBA)^75,77^(Fig. S5h).

#### Compounding consequences of decreased spiking

Importantly, the immediate, direct consequence of any of these defects is a decrease in metabolic spiking that would lead to a decreased dopamine tone at the postsynaptic terminals in striatum (see Supp. Table 3.4 for other co-factors). Here the absence of long-term activation of D2-autoreceptors may also lead to an increased PKA-mediated phosphorylation of tyrosine hydroxylase, and therefore increased dopamine production (Fig. 6b black arrows). Due to lack of metabolic spikes, this dopamine would be minimally transported into the already full synaptic vesicles and linger in the cytosol. Prolonged cytosolic dopamine would lead to a dysregulation cascade^78^ and disruptive aldehyde and quinone production (DOPAL^79^, DA-Q^80^). For example, DOPAL reacts with lysine-rich proteins, such as those in ubiquitin and Alpha-synuclein (*α*-Syn), rendering them obsolete or even deleterious^81,82^ (Fig. S5c, d, f, g). Further DOPAL-stuck ubiquitin itself can disrupt autophagosome formation and may lead to Lewy bodies (Fig. S5c). Such dopamine dysregulation cascades, originating from loss of metabolic spikes, would–over many months or years–degrade SNcDA neurons and lead to their ultimate demise due to increased repair costs of obstructions (increased non-spiking costs).

### VI. Testable predictions and exceptions

We claimed that when ATP usage is lower than baseline levels, neurons must spike or die. We proposed a test to verify such a drastic response in Purkinje cells which naturally exhibit high firing rates. Forcing them to resting membrane potential should reveal ROS that is otherwise compensated by metabolic spikes (Fig. 7a-c), and as such become experimentally observable^83,84^. To establish causality, we also prescribe interventions in the neuron that would decouple mitochondrial metabolism and its excitability (Fig. 7d-f).

Some of these perturbations have already been performed indirectly in independent studies (Supp. Table 3.1). For example, a connection between the metabolism of a neuron and its spiking output can be gleaned in hippocampal pyramidal neurons of mice, where inhibition of DHODH lowers the firing rate set-point^85^. In another study, transient DC hyperpolarizing current which suppressed firing lead to a long lasting increase in excitability^86^.

Some exceptions to our hypothesis may apply. For example, neurons which rely on graded potentials, such as retinal photoreceptor cells, as well as neurons in *Caenorhabditis elegans* which also operate with graded potentials for synaptic transmission^87^, may not rely on spiking for metabolic homeostasis. Interestingly, *C. elegans* prefer low O_2_ environments and display an altered sleeping pattern^88^, suggesting an entirely different path of ROS homeostasis. More generally, any differences in a neuron’s metabolic profile, its developmental stage, neurotransmitter production, its reliance on astrocytes, as well as its relative location with regard to brain region, blood vessels, and ventricles may also play a role in the unique composition of ion channel expression that dictates a neuron’s response. In that vein, neurons that rarely or never experience RETROS conditions may not have compensatory mechanisms and thus may not initiate metabolic spikes.

In addition to neuronal idiosyncrasies, experimental conditions, such as use of growth factors, temperature related changes, general slice health, the composition of pipette solutions and artificial cerebrospinal fluid, and even optogenetic stimulation protocols may affect the homeostatic ion concentrations and thus the metabolic status of the neurons. For example, the use of tetrodotoxin (TTX) to block action potential initiation is a common perturbation, but the effects of interactions between the secondary alcohols of TTX and ROS^89,90^ may complicate the interpretation of such experiments in the framework of our hypothesis.

#### Further implications

Our hypothesis may have some additional implications beyond the supporting models we have presented here. For example, we now know that over their development, neurons undergo a metabolic shift to depend heavily on their mitochondria^33^. At such a key transition time point, metabolic spikes may be involved (i) in priming cell specific intrinsic firing patterns, (ii) in refining the neural circuits and (iii) in establishing a global firing patterns such as those observed in cortical slow waves.

Further, metabolic spikes may also play a role in the onset of some types of epileptic seizures. In our framework, excess run-away activity can be formulated as over-activation of metabolic spikes driven by excess RETROS in neurons or due to hyper-sensitivity of the ion channels sensing the metabolic status. Intriguingly, in some patients where dietary restrictions serve as a therapeutic measure against epilepsy, perhaps the neural metabolism switches in ways that lowers homeostatic metabolic spiking and thus seizure vulnerability.

Finally, one may be tempted to speculate that the evolutionary origin of action potentials (a millisecond binary event) may have been driven–in part–by metabolic pressures. To quickly process multiple signals from external sources (such as graded potentials or peptidergic inputs) neurons may have evolved towards an on-demand ATP production strategy, utilising a lean ATP production cycle with minimal carbon reserve inventory and an increased dependence on ATP-efficient mitochondria. As such, the baseline energy requirements of the cell could be rapidly met by maintaining a healthy mitochondrion population, and additional ATP can be quickly synthesised when needed. The only caveat of this arrangement is that when ATP usage drops below baseline, cells must be able to deal with imminently changing RETROS conditions. Since any upstream compensation (such as e.g., changing the numbers of mitochondria) would happen at slow timescales, only a transient and immediate increase in ATP consumption could avoid RETROS poisoning. Increasing ATP usage by increasing hyperpolarising leak currents is impractical as it would render the cell unresponsive. Instead, producing a fast and reversible depolarisation to shed ATP, possibly at a specialised (axon initial segment-like) site may have been the preferred strategy. Synaptic integration could well have co-opted this mechanism to transmit outputs as digital signals with better reach, providing the evolutionary advantage of rapidly processing multidimensional spatio-temporal inputs.

## Conclusion

In conclusion, metabolic regulation of neural excitability may present a crucial puzzle piece to contextualise the neuronal dynamics that form the basis of all behavior and may have large implications for the evolution of the neural code. If true, metabolic spikes add to the many riches mitochondria bestow upon us.

## Acknowledgements

We thank Prof. C. Nazaret, and Prof. J-P. Mazat for sharing the code of their mitochondrial model. We also thank G. Miesenböck, E. Marder, L. Abbott, A. Kempf, P. Hasenhuetl, W. Podlaski, F. Zenke, E. Agnes, P. Bozelos and G. Christodoulou, and the rest of the Vogels Lab for their feedback.

## Funding

This work was funded by Wellcome Trust and Royal Society Sir Henry Dale Research Fellowship (WT100000), a Wellcome Trust Senior Research Fellowship (214316/Z/18/Z), and a UK Research and Innovation, Biotechnology and Biological Sciences Research Council grant (UKRI-BBSRC BB/N019512/1).

## Author contributions

The ideas in this manuscript were conceived by CC and developed by CC and TPV. All simulations were performed and all figures were generated by CC. Manuscript was written and edited by CC and TPV.

## Competing interests

Nothing to declare.

## Data and code availability

Data sharing not applicable. No datasets were generated or analysed. Code will be made available [here](https://github.com/ccluri/metabolic_spiking) upon publication.

## Methods & Supplementary Material

### 1 Additional discussion

In this work we build a theoretical framework around the hypothesis of metabolic spiking that is based on two main assumptions highlighted in the appropriately coloured boxes, and inspired by circumstantial experimental evidence; they will (hopefully) serve as hypotheses for future experimental studies, some of which we laid out in Figure 7. We explore the validity of our framework in 5 independent models. In the following, we provide additional considerations and experimental evidence for both assumptions and hypothesis, as well as a detailed Methods section that elaborates on the nature of each model’s simplifications. Finally, we discuss the implications of our framework on for a neuro-degenerative disease who’s hallmark symptom is altered spontaneous activity, i.e., Parkinson’s Disease.

#### 1.1 Mitochondrial reactive oxygen species

ATP production in mitochondria routinely leads to the formation of reactive oxygen species (ROS)^1–3^, highly reactive compounds that interfere with cellular processes^4^. Mitochondrial ROS arise when electrons escape from the ETC and react with oxygen to form free radicals. ROS were initially considered as by-products of metabolism that a cell must cope with, however in the last two decades their role in cellular signalling has also gained prominence^5,6^.

ROS is produced at many sites in the mitochondria. In particular, at the mitochondrial complex I, electrons escape from the ETC to produce ROS under two distinct circumstances. During periods of increased energy demands, electrons escape from the flavin mononucleotide sites of the complex I in a process known as “Forward Electron Transport” ROS (FETROS)^7^. In neurons, these conditions could occur, e.g., during periods of high synaptic input, when mitochondria must operate at their maximum capacity (know as “respiratory state 3”^8^). Somewhat counter-intuitively, decreased energy demands can also lead to increased ROS release. When mitochondrial ATP production stalls, e.g., due to scarcity of ADP and concurrent high proton gradient ΔΨ at complex V (respiratory state 4^8,9^), coenzyme-Q molecules carrying electrons reverse their direction of electron transport, jettisoning electrons onto the complex I ubiquinone-binding site, thus allowing them to deviate from the ETC to form ROS^7,10,11^. This phenomenon is called “Reverse Electron Transport” ROS (RETROS) release. While FETROS and RETROS are nearly identical, the circumstances in which they occur are vastly different. FETROS co-occurs with high flux of glucose through the glycolytic pathway, under oxidised ROS scavenger pools, low cytosolic ATP_*C*_ and high cytosolic Ca^2+^ concentrations. Conversely, RETROS co-occurs with low glycolytic flux, reduced scavenger pools, high cytosolic ATP_*C*_ and low cytosolic Ca^2+^ concentrations.

#### 1.2 Role of RETROS in health and disease

There is mounting evidence *in vivo* that ROS and more specifically, RETROS play a key signalling role. For example, it has been demonstrated that RETROS influences longevity and is involved in the coping mechanisms for stress. In fruit flies, RETROS from complex I has been shown to affect lifespan^12^, and increasingly frequent interruptions of electron flow to the ETC^13^ and consequent loss of RETROS has been shown as a hallmark of aging. In another study, exposure to heat stress^14^ has been shown to induce RETROS release which, when suppressed, decreases their survival. A similar effect was also shown in mice with a complex IV knockout^15^. The origin and the magnitude of RETROS formation have also been uncovered in recent years. In homogenates of mouse, rat and human digitonin-treated brain tissue, succinate-dependent H_2_O_2_ generation and the consequent RETROS at complex I potentially contributes nearly 50% of all ROS produced in neurons^16^.

At the behavioural scale, Fernández-Agüera and colleagues have shown that the hyperventilatory response to hypoxia in mice is due to the RETROS signal arising from the complex I of the cartoid body cells. Likewise, Bergmann and Keller^17^ have shown in a mouse *in vitro* study that chemical hypoxia induced by sodium cyanide inhibits complex IV to produce ROS (concurrent with increases in ΔΨ and NADH). These conditions depolarize the motor neurons by increased Na^+^ influx. Kempf et al.^18^ show that the production of RETROS over the course of a day in the dorsal fan-shaped body (dfb) neurons of fruit flies is correlated with their increased firing, and consequent sleep onset. Finally, Dissel and colleagues show^19^ that dfb neurons keep a memory trace of recent metabolic challenges to promote wakefulness, although the pertinence of RETROS was not directly addressed here.

Disruptions to RETROS formation have also been implicated in diseases. In addition to the below mentioned role in Parkinson’s disease, Zhang and colleagues^20^ show that kainic acid induced status epilepticus leads to succinate accumulation that generates RETROS (although it is currently unclear if mitochondiral dysfunction and the resulting ROS are the cause or the consequence of seizures^21^). Similarly, Styr et al.^22^ show that blocking a mitochondrial protein involved in adding electrons to the ETC attenuates seizure susceptibility in Dravet syndrome epilepsy models in mice. Many other studies also implicate the involvement of ROS in neuro-specific sub-fields, such as in neurodegeneration^23^, ischaemia-reperfusion injury^24–26^, in neural development^27^, and synaptic plasticity^28,29^.

#### 1.3 Fate of glucose in neurons

Glucose is the principle energy source for many cells. The glucose that enters the neurons, crosses the blood brain barrier and is absorbed in via insulin-insensitive glucose transporter 3 (GLUT3) transporters. Cellular glycolysis, is regulated by three major enzymes (hexokinase, phosphofructokinase and pyruvate kinase) (Fig. S6a, b). In neurons, the enzyme hexokinase-1 (HK1) irreversibly phosphorylates glucose into glucose 6-phosphate (G6P), effectively locking glucose inside the cell and committing it to be metabolised^30^. Further in neurons, the enzymes phosphofructokinase and pyruvate kinase isozymes M1 are regulated primarily by AMP, ADP, and ATP, with all other regulatory mechanisms nascent^30–35^. Such exclusive control allows for a direct link between glycolysis and a neuron’s metabolic utility. When ATP demand is high, i.e., in FETROS conditions (Fig. S6a), glucose undergoes glycolysis to produce pyruvate to be further metabolised in the TCA. On the other hand, when utility of ATP is low, storing glucose as glycogen – like in many other cell types – is generally not observed in healthy neurons^30,36,37^, and instead the phosphorylated glucose (G6P) maybe processed via the pentose phosphate pathway^38^ (Fig. S6b).

#### 1.4 Pentose phosphate pathway and ROS scavengers

Rerouting glucose via the pentose phosphate pathway (PPP) supports general cellular maintenance through the production of ribose 5-phosphate (R5P) which is used in the synthesis of nucleotides and nucleic acids. These repair pathways are perhaps critical to survival for terminally differentiated post-mitotic cells such as neurons.

More importantly, enhanced PPP has the added benefit of generating reducing equivalents in the form of NADPH (Fig. S6b). NADPH boosts ROS scavenger pools by replenishing the principal redox couples in neurons – glutathione (GSH) and thioredoxin (TXN) (Fig. S6e). Such ROS scavenger pools, through their redox reactions, capture escaped electrons and minimise ROS damage^39^. As NADPH’s availability is considered to be the limiting step of ROS scavenging in cells^40^, any additional pathways that can produce NADPH may also be enhanced if necessary (Fig. S6c).

Under RETROS conditions the scavenger pools operate at their maximum capacity i.e., NADPH produced is fully engaged in scavenging. Consequently, the redox pools persist in their reduced state (Fig. S6e) and any further ROS generated must be quenched by other means due to ROS scavenging overload^39^. Conversely, under FETROS conditions, mechanisms for increased ROS scavenging production are less active, as the pentose pathway is likely suppressed in favour of energy production via glycolysis. Under these conditions, NADPH supply is limited and consequently the redox pools are not replenished and remain in their oxidized state (Fig. S6a, d). Here too, ROS generated must be quenched by other means due to low scavenger availability.

#### 1.5 Other energy sources in neurons

In addition to the standard metabolism of glucose, neurons utilize amino acids, phosphorylated substrates, glycerol, ketone bodies etc., for their energy needs. Another major source is lactate supplied from nearby astrocytes which augments metabolic needs as pyruvate^41^ (see next section). We argue here that, in neurons, the metabolic supply chain to their mitochondria maintains brim-full proton gradients, such that the rate-limiting step of ATP production is pushed to the complex V - the last step of ATP production.

For example, cytosolic pyruvate enters the mitochondrial tricarboxylic acid cycle (TCA, Fig. S6f) where it is converted into acetyl-CoA by un-phosphorylated active pyruvate dehydrogenase (PDH). In neurons, PDH remains unphosphorylated^42^ due to limited expression of its in-activators – pyruvate dehydrogenase kinase (PDK1, PDK3 and PDK4), and high expression of its activator pyruvate dehydrogenase phosphatase (PDP1,2)^30,34^. Cytosolic pyruvate can also form malate^43^ which enters the mitochondria^44^ and can participate in the TCA directly by forming oxaloacetate. In neurons, mitochondrial oxaloacetate either forms alpha-keto glutarate^45,46^ (*α*KG) or combines with acetyl-CoA to form citrate. Notably, under healthy *in vivo* conditions, due to limited phosphoenolpyruvate carboxykinase^47^ (PEPCK) expression oxaloacetate is not turned into PEP. Effectively, this limits the reversal of pyruvate to form glucose and glycogen^36,37,48^. There is also some evidence suggesting that lactate can directly enter the mitochondria to augment pyruvate^49,50^ (but see^51^).

Additionally, glutamate - the most prevalent neurotransmitter and an abundant amino acid, can also be oxidized to *α*KG fuelling the TCA and provide metabolic flexibility^52^. Similarly, acetyle-CoA could be sourced from other metabolic sources such as ketone bodies from the blood, fatty acids and leucine. Irrespective of the metabolic sources, nicotinamide adenine dinucleotide (NADH) and flavin adenine dinucleotide (FADH_2_) produced in the TCA, and other processes such as glycerol 3-phosphate dehydrogenase, electron-transferring flavoprotein (ETF) complex and dihydroorotate dehydrogenase load electrons onto ETC^13^, increasing ΔΨ, keeping the rate limiting step of ATP production to complex V and leading to dearth of ADP, and consequent RETROS as discussed above.

#### 1.6 Lactate and astrocytes

A principle metabolic source in neurons is the lactate produced in the astrocytes nearby^41^. Under heavy metabolic demand in neurons, astrocytes provide auxiliary support by metabolizing glycogen reserves into glucose, which is then used to produce lactate, or by increasing fatty acid oxidation in their peroxisomes to be shuttled to the neurons. Such a cross-talk between astrocytes and neurons is thought it be mediated by the neurotransmitter re-uptake from the extracellular space by astrocytes and the subsequent Ca^2+^ transients in the cytosol of astrocytes. Under low metabolic demands, astrocytes build glycogen reserves^38,53^.

#### 1.7 Battery Analogy

A useful analogy for the role of MS-sensing and metabolic spikes in neurons is that of an electric battery. FETROS conditions arise when the battery is under heavy usage and is also simultaneously being charged. Under these conditions, the battery overheats (akin to ROS production). Spike-prevention provides a chance for the “battery” (the mitochondria) to catch up. RETROS conditions arise when the battery is reaching its maximum charge capacity under light use, but continues to be charged, which also leads to overheating. Metabolic spikes are a mechanism to momentarily use the battery, and thus suspend over-charging. Neurons approaching “nearly-full” battery status must thus spike periodically.

We found the battery analogy useful in the context of the Parkinson’s Disease (PD) and loss of tonic firing in SNcDA neurons. For example, under conditions related to chronic blockade of the complex I ubiquinone binding site (by rotenone) the ‘battery’ is fully chargeable, but the “full” signal is disrupted. Age and sex related PD co-factors are analogous to inefficient batteries that do not reach full charge. Mitochondrial trafficking, tagging, and recycling related faults are analogous to charging dead batteries. In case of ion channel mutations, battery overcharging occurs and the “full” signal is transmitted but the discharge event cannot be initiated. In case of increased per-spike cost, each spike leads to an increased battery discharge, increasing inter-spike interval between battery full states. Finally, faults in lysosomal degradation are analogous to a steady increase in general use of the battery and seldom full states.

### 2 Methods

In this work we begun modelling the effect of metabolism on spiking in a mitochondrial model, which we then reproduced - in the abstract - in a metabolic accounting model, showing that its effects could explain experimentally observed firing properties. We demonstrated that even small changes in specific ion channel properties mediated by metabolism could alter neuronal firing rate dramatically. In the next step, we added metabolic sensing to a network model of integrate-and-fire neurons, reproducing experimentally observed network dynamics. Finally, we explored how two ion channels, –an ATP-sensing potassium channel, and a ROS sensing low-threshold depolarising (sodium and/or calcium) channel– can fulfil the role as metabolic regulators of spiking, and how metabolic dysfunction may lead to symptoms of neural disease.

#### 2.1 Adapted mitochondrial metabolism model

We reproduced and (minimally, see below) adapted a previously published mitochondrial model^54^ that simulates the metabolic components of TCA and ATP production in mitochondria with a limited number of differential equations using mass action kinetics and irreversible reactions. The following section follows closely the appendix A.1 of Nazareth et al., 2009^54^), with a few minor changes. The system evolves as follows:

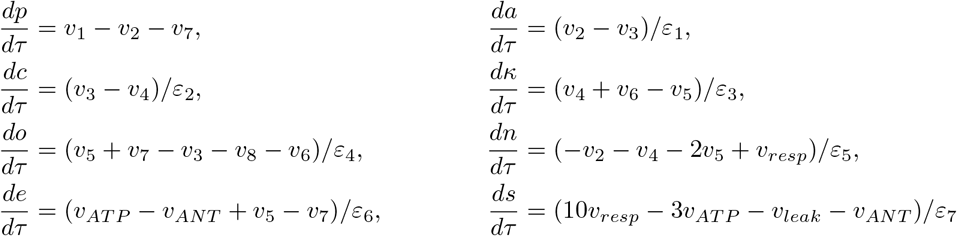

where the constants *ε*_*x*_, *x* ∈ [*p, a, c, κ, o, n, e, s*] are ratios of average product concentrations such that

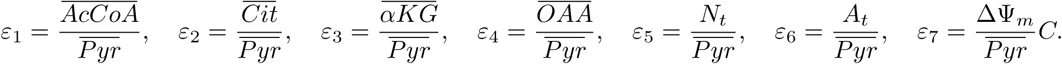

*p, a, c, κ, o, n, e* and *s* are scaled values of pyruvate (*Pyr*), acetyl-CoA (*AcCoA*), citrate (*Cit*), *α*-ketoglutarate (*αKG*), oxaloacetate (*OAA*), NAD+ (*NAD*), ATP (*ATP*) and proton gradient (ΔΨ_*u*_). Their physical values can then be calculated from

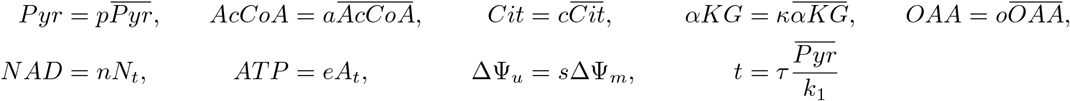

with average values as observed in experiments, i.e., 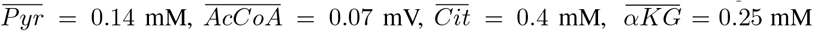 and 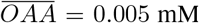. At these substrate concentrations, ΔΨ_*m*_, 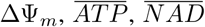 were measured to be 150 mV, 3.23 mM and 0.94 mM. Additionally, *N*_*t*_ = 1.07 mM and *A*_*t*_ = 4.16 mM are the maximum possible concentrations NAD+ and ATP can reach. *τ* is the scaled time, *C* = 6.75e-6 M/V is the scaled capacitance, and *k*_1_ is the rate constant of pyruvate intake to the model (see below).

Reaction rates *υ*_*x*_, *x* ∈ [1, 2, 3, 4, 5, 6, 7, 8, *ANT, ATP, leak, resp*] were given by

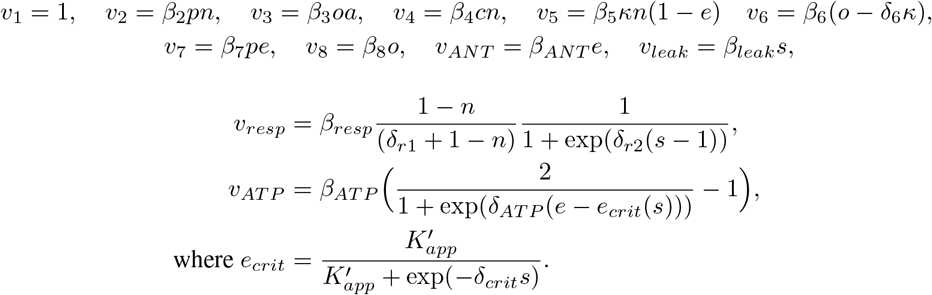

The constants *β* and *δ* are

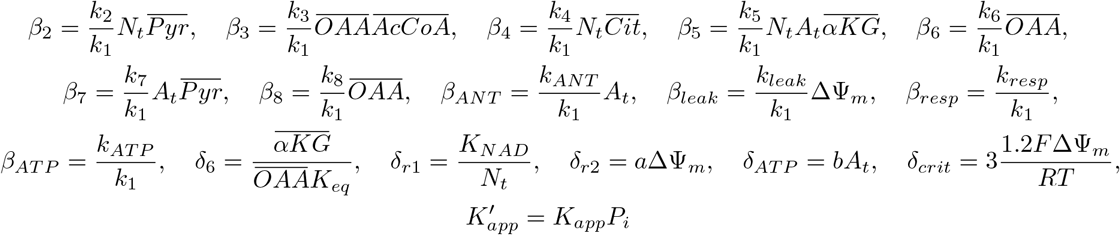

Reaction rates were recorded as 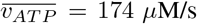 and 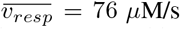. Based on these reaction rates, and on the observed equilibrium concentrations of aspartate and glutamate, additional equilibrium constants were set as *K*_*app*_ = 4.6e-6, *K*_*NAD*_ = 2 mM, *a* = 100 /V, *b* = 4 /V and *K*_*eq*_ = 0.397. *P*_*i*_ = 2.44 mM is the concentration of phosphate ions, F=96485 C/mol is the Faraday constant, R=8.314 J/(molK) is the gas constant, and T=298 K the temperature in Kelvin, equivalent to 24.8 °C.

In order to obtain a steady state metabolism in the model that is consistent with the experimental literature, i.e., respiratory state 3.5, Nazaret et al. estimated *k*_1_ and *k*_3_ as 38 *μ*M/s and 57, 142 /(Ms), respectively. All other rate constants then follow as *k*_2_ = 152 /(Ms), *k*_4_ = 53 /(Ms), *k*_5_ = 82361 /(M^2^s), *k*_6_ = 9.0032 /s, *k*_7_ = 40 /(Ms), *k*_8_ = 3.6 /s, *k*_*resp*_ = 2.5 mM/s, *k*_*ATP*_ = 131.9 mM/s, *k*_*leak*_ = 0.426 mM/(Vs) and *k*_*ANT*_ = 0.05387 /s. In the following, we kept all these rate constants fixed, with the exception of *k*_*ANT*_ and *k*_*leak*_ where noted.

In Nazareth model, *e*, the unitless ATP concentration variable, and *n*, the unitless NAD concentration variable could assume values ∈ (0,1). All other variables, *p, a, c, κ, o* and *s* ∈ [0,∞). Diverging from the Nazaret model, we convert the unitless variable *s* for the proton gradient to ΔΨ, such that

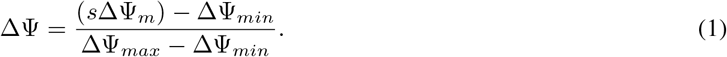

with ΔΨ_*min*_ = 125 mV (for *k*_*ANT*_ >1 /s) and ΔΨ_*max*_ = 190 mV (for *k*_*ANT*_ <10^−3^ /s). Assuming that mitochondria operate in the range between these two extreme metabolic states, we can constrain ΔΨ to values ∈ (0, 1). For clarity, we refer to the unitless variable *e* for mitochondrial ATP concentration as *ATP*_*M*_ (*ATP*_*M*_ = *e*), and we express *k*_*ANT*_ in the units of 10^−3^/s or /ks in the text.

###### Simplifications in the Nazareth model - (I)

The mitochondrial model is a simplified TCA and ATP production model based on mass action kinetics. It is constrained by experimentally observed steady state values of consumption and production in isolated mitochondria. Importantly, the model does not include pyruvate production from glycolysis or lactate. **(II)** The model also forgoes AMP production, or de-novo synthesis. **(III)** Finally, creatine phosphorylation, uncoupler proteins, mitochondrial fusion, fission, mitochondrial migration, mitochondrial permeability transition pore opening and mitophagy are not modelled. These simplifications allow for a focus on the crucial aspects of metabolism in neurons that makes the model by Nazareth et al. a good starting point to explore interactions of metabolism and neural excitability.

#### 2.2 Integrating mitochondrial metabolism model into a neuron model

To combine mitochondiral ATP production with neural ATP consumption we added a cytosolic ATP-consuming process –modelling e.g. Na-K pumps– such that the concentration of [ATP_C_] follows 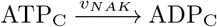, where the rate of the reaction *υ*_*NAK*_ = *k*_effective_[*ATP*_*C*_], with *k*_effective_ as an abstract function that depends on [Na^+^], [K^+^] and other quantities. For simplicity, we considered mitochondria to be the only ATP source^†^ such that ATP_M_ = ATP_C_ and *k*_effective_ = *k*_*ANT*_. To model calcium influx into the mitochondria after a given spike, adjust *k*_*leak*_, effectively modulating the leak of protons from the intermembrane space into the matrix 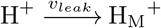, where *υ*_*leak*_ = *k*_*leak*_ ΔΨ. Note that Ca^2+^ in the matrix doesn’t change any other reaction rates^††^. We consider its efflux to the cytosol is much slower than the spike related transients, and thus absorbed efflux in *k*_*ANT*_ (see below).

###### Assumption I

In neurons, the rate limiting step of ATP production is located at its last step, i.e., complex V (ATP-synthase) of the electron transport chain in their mitochondria.

###### Simplifications in the metabolism coupled neuron model

Without loss of generality we can make the following simplifications:**(I)** The mitochondria are the only ATP source and the ATP concentrations in a neuron’s cytosol and in its mitochondrial matrix are at equilibrium. **(II)** The Ca^2+^ that enters the mitochondria after each spike is modelled directly as a decrease in ΔΨ. Ca^2+^ buffering in the mitochondria is not modelled explicitly and its eventual extrusion from the matrix is implicitly included in the non-spiking costs. Due to lack of neuron specific data, the regulation of ATP production by free Ca^2+^ in the mitochondria is not included in our model.

#### 2.3 Modelling the effect of a spike

The metabolic expense of a spike was modelled as a rapid transient increase of the effective ATP consumption rate constant *k*_effective_ (= *k*_*ANT*_), followed by slow decay back to baseline, such that *k*_*ANT,spike*_(*t*) = *k*_*ANT*_ + *Q*_*T*_ (*f*_*Q*_, *t*), where *k*_*ANT*_ is thus the non-spiking related costs that can be assumed to vary very slowly compared to spike related transients. Specific examples of non-spiking related cost values are indicated as ⋆ symbols in all the figures. *Q*_*T*_ (*f*_*Q*_, *t*) is the per-spike cost that occurs at time T (the instance of *V > V*_*θ*_). We model *Q*_*T*_ (*f*_*Q*_, *t*) such that 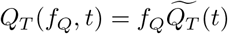, where 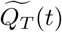 is an exponentially rising (*τ*_*Qrise*_=0.6 ms) and decaying (*τ*_*Qfall*_=100 ms) function with unit maximum (at *t*_*lag*_), i.e.,

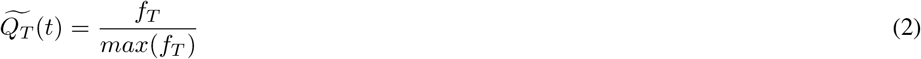

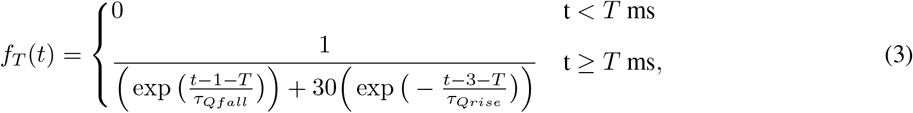

These costs were assumed to include all the metabolic costs incurred after spike initiation, i.e., spike generation, neurotransmitter release, re-uptake and synaptic vesicle loading. Depending on the modelled neuron type, spike costs vary, which we include as multiplicative factor *f*_*Q*_ that has the units of *k*_*ANT*_ (1/ks). Under wild-type conditions, i.e., assuming a steady mitochondrion distribution at the spike initiation micro domains, *f*_*Q*_ is fixed^§^ for a given neuron. As mentioned above, each spike also initiates Ca_2_^+^ entry into the mitochondria^56^. As a divalent cation, Ca_2_^+^ decreases ΔΨ, which we model as a transient increase in leak *k*_*leak*_. For simplicity, recovery of *k*_*leak*_ follows the same time course as *Q*_*T*_ (*t*) with a multiplicative factor *f*_*MCU*_, such that *k*_*leak,Ca*_(*t*) = *k*_*leak*_ + *f*_*MCU*_ *Q*_*T*_ (*f*_*Q*_, *t*). Unless otherwise mentioned, *f*_*MCU*_ is set to 0.01 M/V. For example, for *f*_*Q*_= 1/ks, post-spike calcium entry to the mitochondria increases *k*_*leak*_ from 0.426 mM/s to a maximum of 0.436 mM/s (At *t*_*lag*_ from onset) for default *f*_*MCU*_ .

###### Assumption for modelling the effect of a spike - (I)

There is a transient increase in the cytosolic metabolic cost after each spike (per-spike cost Q). **(II)** The non-spiking metabolic costs vary slower than the per-spike costs.

###### Simplification in modelling effect of a spike

Without loss of generality we can make the following simplifications: Ca^2+^ entering the mitochondria follows the same time course as the per-spike costs (Q).

#### 2.4 Calculating ROS levels

We model ROS levels based on the “redox-optimized ROS balance hypothesis”^39^, stating that ROS levels are a function of a cells’s redox potential and follow a non-monotonic V-shaped curve (Fig. 1j). The cell’s redox can be deduced from the ratios of the so-called redox couples NADH/NAD+, CoQH_2_/CoQ, GSH/GSSH, NADPH/NADP+, and their respective half-cell reduction potentials. Minimum ROS levels are achieved when all redox couples are balanced, i.e., their ratios approach 1. ROS levels increase at the redox potential extremes, i.e., when the ROS couples approach either the fully reduced or the fully oxidized state.

The oxidised state may be approached due to increased traffic on the ETC during high ATP demands, when the availability of mitochondrial substrates (pyruvate, NADH and FADH_2_) becomes rate-limiting. The redox couples are oxidized (i.e. the denominators become bigger than the numerators) and cannot be re-balanced because energy production is prioritised over ROS scavenging, i.e., glucose is routed via glycolysis to pyruvate production instead of the pentose phosphate pathway, hence limiting the scavenging capacity due to low cytosolic NADPH (see below). In other words, ROS production is relatively stable, but scavenging capacity decreases.

The reduced state may be approached when the ETC stalls and ADP in the mitochondria is the rate-limiting step. Here, the redox couples are reduced (i.e. the denominators become smaller than the numerators) and scavengers operate at their maximum capacity. Glucose is routed to the pentose phosphate pathway instead of glycolysis, consequently cytosolic NADPH production and its utility as a scavenger are at their maximum. In other words, ROS production exceeds the maximum scavenging capacity.

We can express these biochemical relations in mathematical terms as

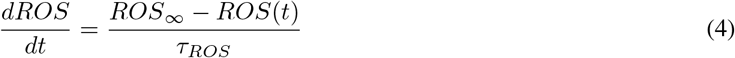

whose steady state *ROS*_∞_ is determined by *ATP*_*M*_ and ΔΨ values with a cubic relationship.

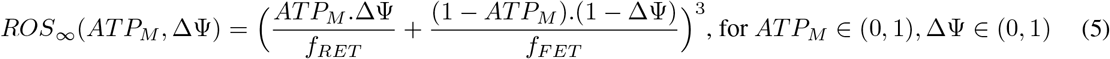

FETROS and RETROS conditions are thus coupled to the rate at which ATP is produced in the mitochondria as well as to the rate at which ATP is consumed by cellular processes. Under heavy metabolic demands the ATP consumption is high and any ΔΨ is depleted to produce ATP (FETROS). Under low metabolic demands both ΔΨ and ATP are high (RETROS). *f*_*RET*_ ∈ (0, ∞) and *f*_*FET*_ ∈ (0, ∞) are tuning parameters that determine the relative amplitudes of RETROS and FETROS production. Unless otherwise mentioned they are set to 1.

Additionally ROS dynamics are characterised by a time constant (*τ*_*ROS*_) which depends on ATP consumption and peaks near ROS minimum such that

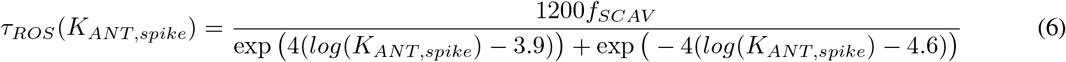

Here, *f*_*SCAV*_ is a multiplicative factor which modulates the intensity of scavenging in a neuron. Unless otherwise mentioned we set *f*_*SCAV*_ to 1ms. We also tested conditions when *τ*_*ROS*_ was kept constant and independent of ATP consumption.

###### Assumption in the ROS production model

Neurons, depending on their physiological operational conditions, can face a range of metabolic demands and consequently may be exposed to non-monotonically varying ROS levels. At ideal metabolic preparedness, ROS levels are minimal and increase both when higher (FETROS) and lower (RETROS) energy demand occurs.

###### Simplifications in the ROS production model - (I)

To express the non-monotonic nature of the ROS levels, we chose a cubic function of normalized ATP and ΔΨ values (Eq.5). In real neurons, these steady state values could be determined by measuring ROS levels while suppressing firing. **(II)** The time constant of the ROS levels serves as a proxy for the ROS scavenger response of the neuron.

#### 2.5 Integrating metabolic signal into an accounting model

To explore the hypothesis of ROS homeostasis quantitatively, we first implemented a metabolic accounting model which changes its metabolic signal (MS) according to FETROS and RETROS events such that

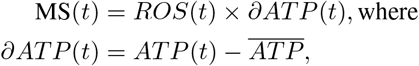

with 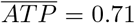 as the steady state *ATP* at minimum ROS. We define two thresholds, *θ*_*RET*_ and *θ*_*FET*_ such that when MS*> θ*_*RET*_ the neuron initiates a metabolic spike and enters a refractory period *t*_*ref*_. When MS*< θ*_*FET*_, even synaptically driven spikes cannot be initiated. Two parameters thus dictate the intrinsic firing property landscape: (i) the minimum inter-spike interval between successive spikes *t*_*ref*_, and (ii) the lag time *t*_*lag*_ after a spike until the ATP costs affect MS (e.g. related to Na-K pump activation). *t*_*lag*_ is defined as the time between spike initiation *T* and the peak of *Q*_*T*_ (*t*), and can be varied by changing *τ*_*Qrise*_ (eq. 3).

###### Simplification in accounting model

The metabolic signal (MS) is modelled as the product of ROS levels and ∂ATP. MS is a sufficiently unambiguous signal for the ion channels to gauge whether the neuron is in FETROS or RETROS regime. In reality multiple ion channels may respond to a metabolic signal (For example, Ca^2+^ or cAMP levels) or to a combination of several metabolic signals (For example, ATP and H_2_O_2_ levels) that a neuron experiences.

#### 2.6 Dorsal fan shaped body neuron model

We modelled the competitive effects between the A-type and non A-type K channels on spiking in dfb neurons in a single compartment model with a single sodium channel type, as well as a delayed rectifier and an A-type potassium channel model. The membrane potential thus follows

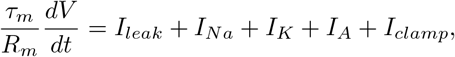

where leak, sodium, delayed rectifier and A-Type potassium currents can be calculated as

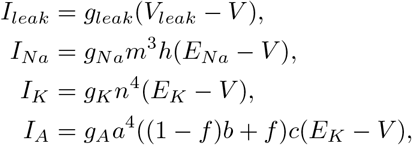

respectively, and *I*_*clamp*_ is the additional externally injected current. Here *f* is the probability of hyperkinetic *β* subunits filled with NADP+ (at sleep onset) instead of NADPH (post-sleep). Parameter values *V*_*leak*_ =-40 mV, *τ*_*m*_ =100 ms, *R*_*m*_ = 1 GΩ, *E*_*K*_ = -60 mV, *E*_*Na*_ =40 mV, *g*_*leak*_ =1 nS, *g*_*Na*_ =1200 nS, *g*_*K*_ =90 nS, *g*_*A*_ =80 nS are based on typical values^57,58^. The gating variables *m, h, n, a, b*, and *c* are standard variables calculated as

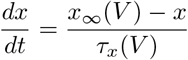

where x ∈ (*m, h, n, a, b, c*). *m,h* and *n* are modified Hodgkin and Huxley variables^57^ for Na and K_DR_ channels; the K_*A*_ variables *a, b* and *c* are based on ion channel studies in ventral cochlear nucleus neurons^58^. In detail, the tuned variables present themselves as

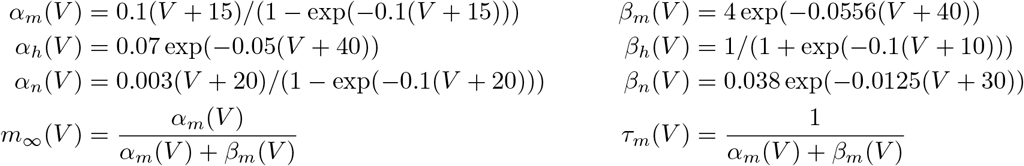

for *m,h*,and *n*, and

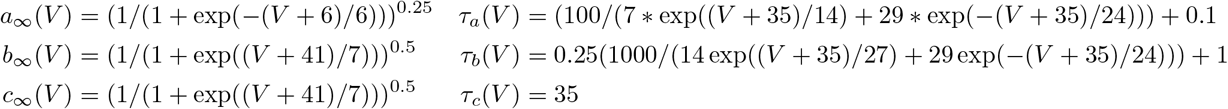

for *a,b*, and *c*. Here *a* is the fast activating gate, *b* is fast inactivating gate and *c* is a slow inactivating gate. Near sleep conditions are modelled as *f* =0.7. The contribution of the inactivating gate *b* to the A-type current is less pronounced and thus the channel remains largely non-inactivating. We model awake conditions as *f* =0 when the *b* gate rapidly inactivates the A-type current.

Due to lack of relevant data for the gating variables, ion channel conductances and morphology of this cell, we hand-tuned the gating variables and conductances to match experimental observations^18^. We limited manipulations of these ion channels to changes in the inactivation gating variable, but other mechanisms and other ion channels may also be involved in modulating firing in these cells.

###### Assumption in dFB neuron model

We assume, as in the experiment, that the changes in excitation are only due to changes in one ion channel.

###### Simplification in dFB neuron model

Dorsal fan shaped body neuron is modelled as a point neuron with three ion channels. Their morphology, channel distribution, and channel properties are not well characterised.

#### 2.7 Recurrent network model

To study the effects of metabolic spiking at the network level we constructed a network of 10,000 leaky integrate-and-fire neurons^59^. Each integrate-and-fire neuron is characterized by a time constant, *τ* = 20 ms, and a resting membrane potential, *V*_*rest*_=-60 mV, and a spike threshold, *V*_*θ*_=-50 mV. After each spike, the membrane potential is clamped at *V*_*rest*_ for a refractory period of *t*_*ref*_ (see below). To set the scale for currents and conductances in the model, we use a membrane resistance of 100 MΩ. In addition to the standard membrane currents (leak, excitatory and inhibitory synaptic currents) we added a metabolic current, such that the membrane voltages are calculated as.

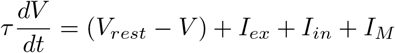

where

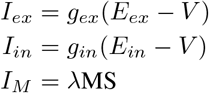

Reversal potentials are *E*_*ex*_ = 0 mV and *E*_*in*_ = −80 mV. The synaptic conductances *g*_*ex*_ and *g*_*in*_ are expressed in units of the resting membrane conductance. Neurons in the network are either excitatory or inhibitory with 4:1 ratio and 2% connectivity. When a neuron fires, the appropriate synaptic variable of its postsynaptic targets are increased, *g*_*ex*_ → *g*_*ex*_ + Δ*g*_*ex*_ and *g*_*in*_ → *g*_*in*_ + Δ*g*_*in*_ for an excitatory and inhibitory presynaptic neuron, respectively. Otherwise, these parameters obey the following equations:

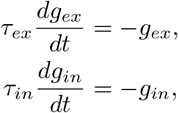

respectively, with *τ*_*ex*_ = 5 ms and *τ*_*in*_ = 10 ms. *I*_*M*_, the metabolic current, emulates the combined effect of several ion channels as a current injection of *λ* that is controlled by the cumulative metabolic signal MS. Here *λ*, was set to 25 mV. In the absence of an explicit mitochondrial module in our model, MS serves as a proxy for current *ATP* consumption rate,

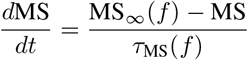

MS largely depends on the steady state metabolic consumption MS_∞_ due to synaptic inputs *f*, as well as output spikes (see below), such that

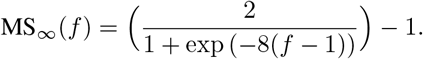

The temporal dynamics of MS are governed by a characteristic time constant *τ*_*MS*_

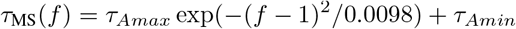

with *τ*_*Amax*_ = 1000 ms and *τ*_*Amin*_ = 300 ms which both also depend on *f* which is expressed as the fraction of synaptic inputs relative to its metabolic optimum *L*, as follows

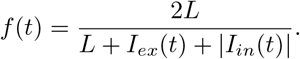

The optimal metabolic load number *L* refers to the point of minimum ROS production in the earlier mitochondrial model above, and is used as a neuron-specific, predefined value from a standard normal distribution with mean *μ* = 2 nA, and standard deviation *σ* = 0.5 nA. |*I*_*in*_| is the absolute value of *I*_*in*_. When *I*_*ex*_ + |*I*_*in*_| = *L, f* (*t*) = 1, and therefore MS_∞_(*f*) = 0, and there is no additional metabolic current (*I*_*M*_ = 0). When a neuron receives less input than its metabolic optimum *L*, i.e., *I*_*ex*_ + |*I*_*in*_| *< L*, then *f* (*t*) *>* 1 and MS_∞_(*f*) *>* 0, and therefore *I*_*M*_ *>* 0 – the metabolic current is depolarizing. Conversely, when the neuron receives much larger input than its optimum, i.e., *I*_*ex*_ + |*I*_*in*_| *> L, f* (*t*) *<* 1, MS_∞_(*f*) *<* 0, and therefore *I*_*M*_ *<* 0 – the metabolic current becomes hyperpolarizing.

In addition to synaptic inputs, MS is also affected by spikes. To model this effect in integrate and fire neurons, MS is decreased such that MS = MS − *q* after each spike, where q is 0.1 and *t*_*ref*_ = *t*_*def*_ − *α*MS, where *t*_*def*_ is 5 ms and *α* is 3 ms and captures changes to spike-adaptation.

###### Simplification in recurrent neural network model

The metabolic expense of excitatory and inhibitory currents are modelled as equal. In reality, metabolic cost of these currents will depend on the specific composition of the involved ion channels (AMPA vs. NMDA, GABA_*A*_ vs. GABA_*B*_).

#### 2.8 SNcDA neuron model

We constructed a leaky integrate and fire neuron coupled to the Nazaret mitochondrial model^54^, and with an ATP-sensing potassium channel and a ROS-sensing low-threshold depolarising (sodium and/or calcium) channel as a model of a SNcDA neuron. We set the non-spiking ATP expenditure to 25 /ks and per-spike cost Q to 30 /ks. At this baseline expenditure and in the absence of input, the neuron elicits tonic metabolic spikes at a frequency of 4 Hz. Similarly to above, the equations that govern neuronal dynamics are given by

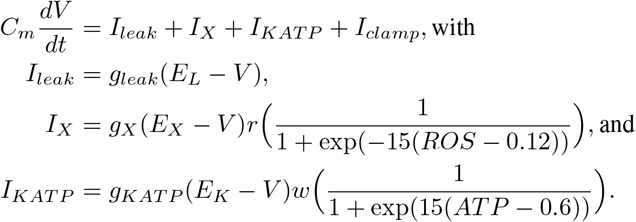

*E*_*L*_, *E*_*X*_ and *E*_*K*_ are -68 mV, 40 mV, and -70 mV respectively. *g*_*leak*_, *g*_*X*_ and *g*_*KATP*_ are 15 nS, 110 nS and 100 nS respectively. Here, *V* is the membrane potential and *C*_*m*_ = 100 pF is the membrane capacitance. *I*_*clamp*_ ∈ (0, 0.3 nA) is a stimulating current that can be adjusted to drive the neuron to spike. *I*_*leak*_ is the leak current; *I*_*X*_ and *I*_*KATP*_ are depolarizing and hyperpolarizing currents that respond to the ROS and low-ATP respectively. The variable *ROS* is calculated from ΔΨ and *ATP* (Eq.5), which in turn are obtained from the adjusted mitochondrial model (see 1.4) that is coupled to the neuronal dynamics through *K*_*ANT,spike*_ and *K*_*leak,Ca*_ (Section 1.3). The gating variables for *I*_*X*_ and *I*_*KATP*_, r and w are given by:

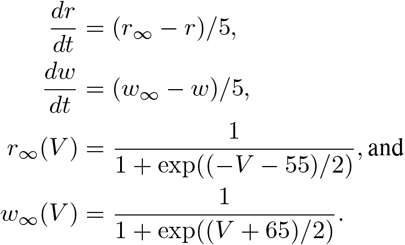

When *V* reaches spike threshold, *V*_*θ*_=-40 mV, a spike is initiated and the membrane potential is set back to at *V*_*rest*_, and *r* is set to 0 to emulate channel inactivation.

We alter the model to emulate specific disease states, i.e., RETROS collapse as a consequence of chronic rotenone exposure, increases in per-spike cost, and an overall increase in non-spike related metabolic expenditure. Chronic exposure to rotenone alters mitochondrial ROS production such that RETROS is diminished, and FETROS is enchanced^*^. We can capture this in our model by setting *f*_*RET*_ = 1.25 and *f*_*FET*_ = 0.8 in eq.5. Note that to model changes related to age, sex or poor mitochondrial trafficking only the RETROS production, i.e., *f*_*RET*_, must be changed. Increases in per-spike cost are achieved by increasing Q by three fold relative to control such that Q=90 /ks. An overall increase in non-spike related metabolic expenditure is achieved by setting the baseline expenditure to 60 /ks (golden star in Fig. 4).

###### Simplification in SNcDA neuron model

Only two metabolic state sensing ion channels are modelled explicitly. These channels represent a hypothetical ROS sensing depolarizing channel and an ATP sensing hyperpolarizing channel.

#### 2.9 Software and figures

Simulations were carried out in python^62^ using numpy^63^, brian^64^ and powerlaw^65^ packages. The human brain sagittal section was obtained from wikipedia (illustrated by Patrick J. Lynch). The neuron drawn on it was based on reconstructions of rat SNcDA neuron^66^. Purkinje cell morphology (ID: NMO_10074) was taken from neuromorpho^67^. The figures were produced in matplotlib^68^ and compiled in libredraw.

### 3 Supplementary Tables

#### 3.1 Experimental observations in agreement with our hypothesis

**Table 1:** Experimental observations in agreement with the intrinsic excitability hypothesis.

| *No.* | *Perturbation* | *Effect* | *Reference* |
| --- | --- | --- | --- |
| 1 | miniSOG/AOX | Genetically-encodable light-induced stimulation of mini singlet oxygen generator (miniSOG) adds cytosolic ROS and increases firing rate. Conversely, expressing plant mitochondrial alternative oxidase (AOX) lowers CoQH<sub>2</sub> pools in ETC and decreases the firing rate of dorsal fan-shaped body neurons in fruit fly, changing their sleep state | 18. |
| 2 | DHODH | Mitochondrial protein dihydroorotate dehydrogenase (DHODH) involved in de novo pyrimidine biosynthesis also increases CoQH<sub>2</sub> pools in the ETC. Inhibition this lowers the firing rate set-point in CA1 pyramidal neurons. | 22 |
| 3 | Low lactate | Lactate dehydrogenase in neurons produces pyruvate from lactate. It's inhibition is reported to lower excitability in neurons and suppressed seizures in an in vivo mouse model for epilepsy. | 69 |
| 4 | Pipette | Increased excitability of neurons in slice compared to in vivo seen in layer 2/3 pyramidal neurons. This is potentially due to decrease in the long range network connectivity and therefore decrease in non-spiking costs. | 70 |
| 5 | DC hyperpolarization | Prolonged DC current injection to inhibit spontaneous firing induces long-lasting increase in excitability. Prohibiting neurons to fire can increase ETC stalling (more CoQH<sub>2</sub> pools) and may enhance the homeostatic metabolic spiking. | 71 |
| 6 | Creatine deficiency | Mutations in the SLC6A8 gene can cause X-linked creatine transporter deficiency with its hallmark symptoms including epilepsy. Lack of creatine in neurons would lead to lowered ATP buffering and may prompt increases in metabolic spiking and thus vulnerability to seizure onsets. | 72 |
| 7 | Low-carb diet | Diets low in carbohydrates have long been prescribed as counter measures to some drug-resistant epilepsy's. Here, neurons may rely more on differently regulated ketone bodies and amino acids for their metabolic needs and decrease their dependence on pyruvate. Perhaps this changes the neuronal metabolism in ways that lead to fewer metabolic spikes and thus lower seizure onset | 73,74 |

#### 3.2 List of potential ion channels that may sense metabolic signals

**Table 2:**
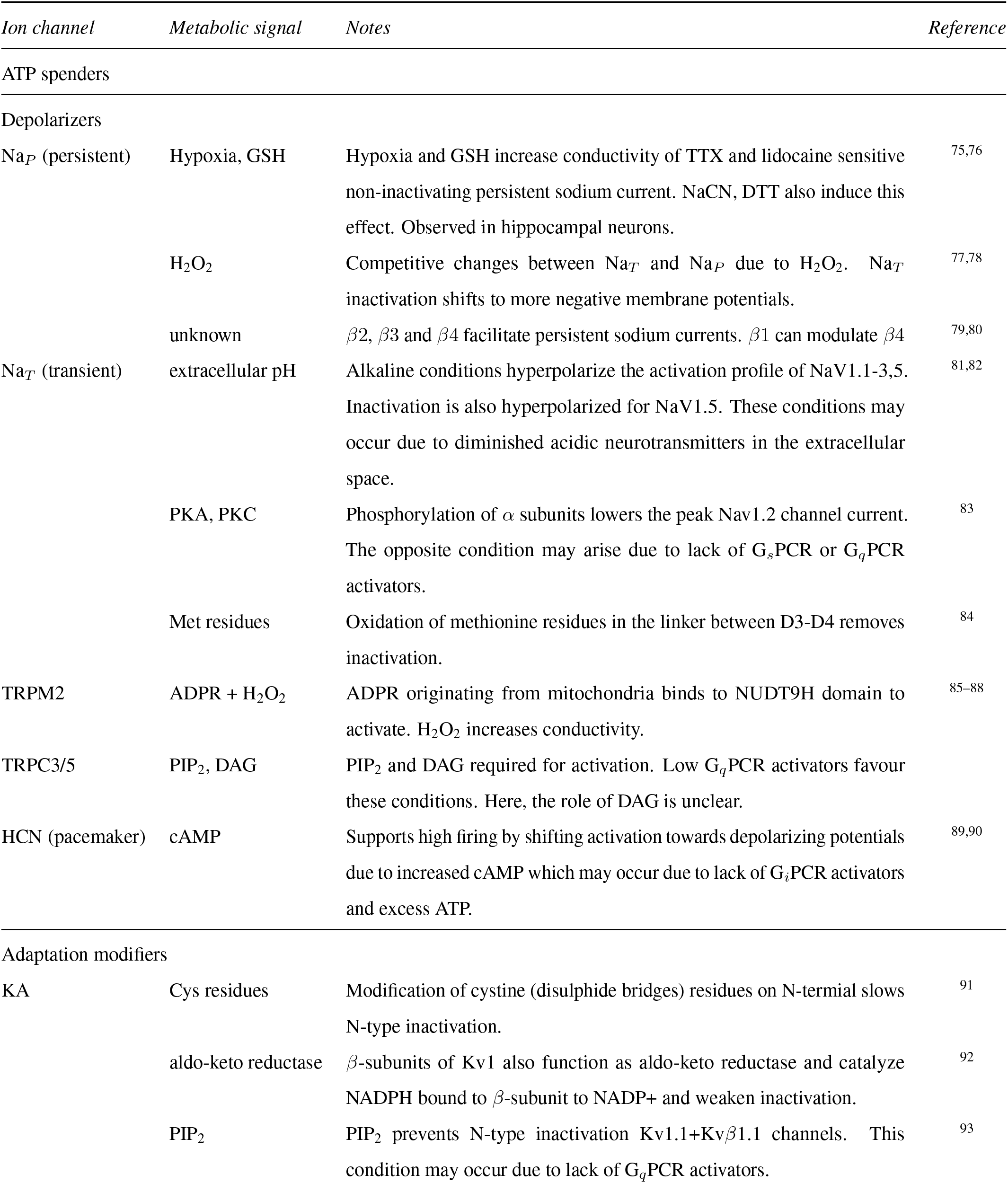
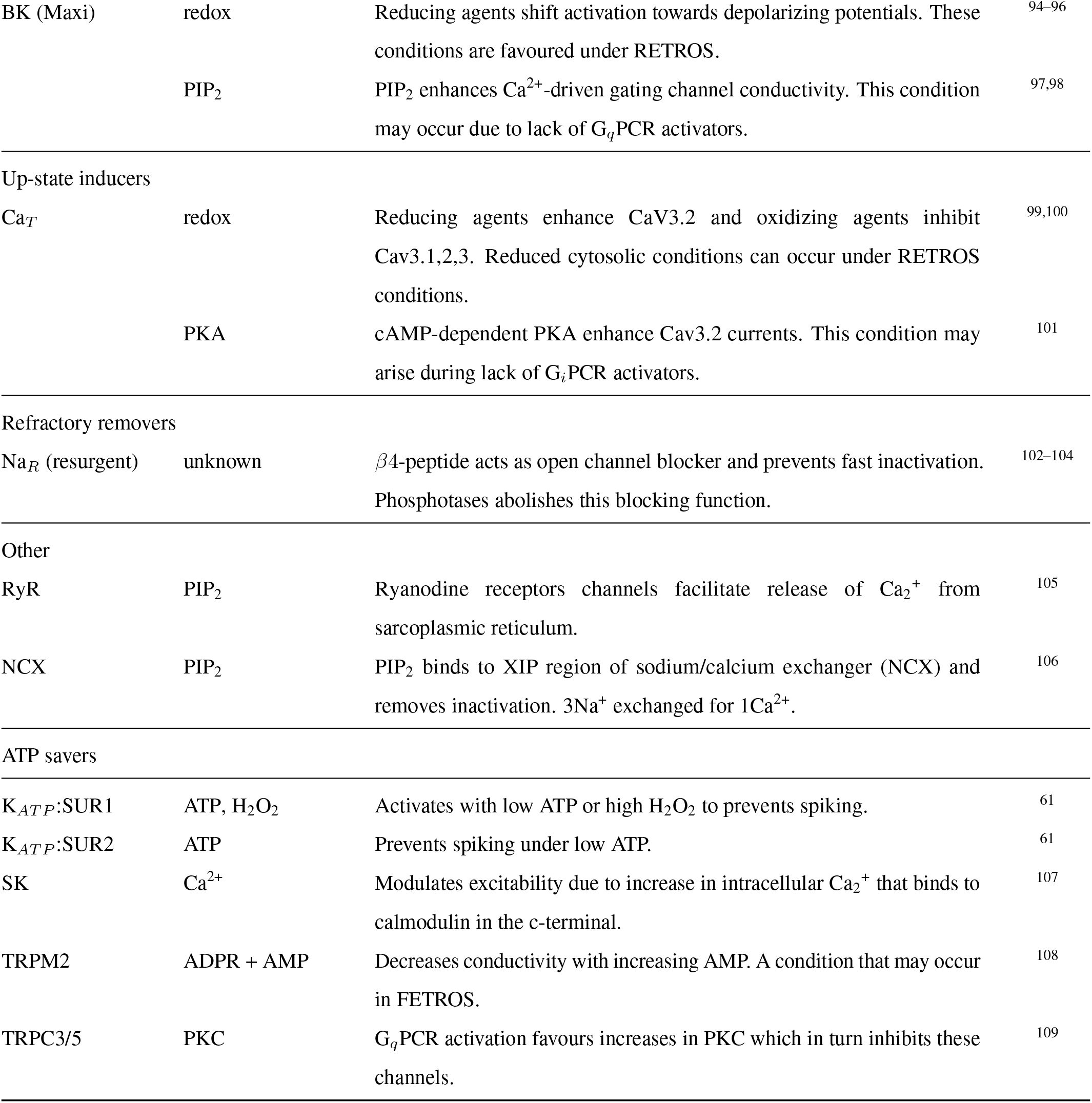
Potential ion channels and their corresponding metabolic signals that align with the intrinsic excitability hypothesis.

#### 3.3 List of co-factors involved in Parkinson’s Disease

**Table 3:** Co-factors involved in Parkinson’s. Co-factors such as *α*Syn, LRRK2 and HSC70 play multiple roles.

| *Co-factor* | *Gene/Intervention* | *Effect* | *Reference* |
| --- | --- | --- | --- |
| Environmental | Chronic rotenone/MPP+ | Competitive binding at Q-binding site of complex I | 7 |
| Age | LDH1/5 | Drop in ratio of LDH1 to LDH5 with age and lower puruvate | 110,111 |
|  | ETC/TCA | Inefficiencies in mitochondrial energy production | 111,112 |
| Sex | X-linked | Vulnerability of genes expressed on X-chromosome e.g.,<br>NDUFA1/NDUFB11 encoding for complex I sub-units, RAB39B | 113,114 |
| Mitophagy | PINK1/PARKIN | Defects in tagging and trafficking of degraded mitochondria | 115,116 |
|  | Ubiquitination | Faults in ubiquitination and repair through deubiquitination<br>(USP30/35) | 117,118 |
| Myelination | Aberrations or loss | Increased sodium, potassium leaks in axon | 119 |
| Vesicle | V-ATPase, VMAT2 | Defects in loading vesicles with dopamine | 120–123 |
| Exocytosis | Spike ($\text{Ca}^{2+}$) | $\alpha\text{Syn}$, HSC70, GEF, Rab-GTPases, SNARE proteins | 122–125 |
| Re-uptake | SLC6A3 (DAT) | Faults in dopamine active transporter (DAT) | 126,127 |
| Endosome | Clathrin coating | Dynamin, Auxilin, Endophilin, Rab-GTPases, $\alpha\text{Syn}$ | 120,122,123 |
|  | Clathrin uncoating | HSC70, synaptojanin, Rab-GTPases, RME-8 | 120,122,123 |
| Lysosomal | ATP13A2, ATP6AP2 | ATPase cation, lysosomal accessory protein transporter | 128–130 |
|  | VPS35 | Trafficking between endosome, lysosome and golgi | 131 |
|  | VPS13C | Lipid transport between endoplasmic reticulum and organelles | 132 |
|  | RAB29 | Recruitment and activation of LRRK2 | 133 |
|  | GBA | Lower glucocerebrosidase (GBA) enzyme activity | 130,134 |
|  | Microautophagy/CMA | Faults in HSC70 dependent identification of KFERQ-like motifs<br>for microautophagy and co-chaperone mediated autophagy<br>(CMA) via LAMP2 | 135 |
| $\alpha\text{Syn}$ | SCNA | Curvature sensing protein involved in endosome formation,<br>exocytosis, lysosomes. | 120,123 |
| LRRK2 | Kinase, GTPase | Involved in regulation of signal transduction via<br>RAB-phosphorylation. Interactions with mitophagy(parkin),<br>lysosomes (GBA), vesicular trafficking (RAB-GTPase) and<br>$\alpha\text{Syn}$. | 136–140 |
| HSC70 | Chaperones | Involved in proteostasis including folding and clearance | 124,141 |

#### 3.4 Other potential ROS relieving mechanisms in neurons

**Table 4:** In addition to modulating intrinsic excitability in neurons, other potential ROS relieving mechanisms that maybe enhanced during RETROS or FETROS conditions.

|  | *Process* | *Notes* | *Reference* |
| --- | --- | --- | --- |
|  | RETROS |  |  |
| Pre complex V | Pentose phosphate pathway | Up-regulation and consequent ribose 5-phosphate production may promote repair pathways and protein synthesis. | 142,143 |
|  | Glycogen synthesis | Glucose is stored as glycogen in astrocytes instead of glycolysis. | 144,145 |
| | Uncoupler proteins | Action of UCP2, UCP4 and UCP5 in the mitochondria, can decrease $\Delta\Psi$. However, they also import fatty acids and amino acids and despite their action the pool of CoQH<sub>2</sub> on ETC remains unchanged. | 146–148 |
| Post complex V | Creatine phosphorylation | Phosphorylation of creatine uses ATP to produce phosphocreatine. | 149 |
|  | Lysosomal activity | Favourable conditions for vacuolar-type ATPase-dependent lysosomal acidification. | 150 |
|  | Synaptic changes | Spine changes, vesicle formation and neurotransmitter synthesis. | 123 |
|  | Ca<sup>2+</sup> clearance | Ca<sup>2+</sup> buffered in organelle may be cleared from the cell. | 106 |
|  | Mitochondrial bio-genesis | Favourable conditions for mitochondrial fission and fusion. | 151,152 |
|  | Cellular cargo transport | Promotion of cytoskeletal motor proteins (dynein and kinesin). | 153 |
|  | FETROS |  |  |
| Pre complex V | Glycolysis up regulation | PFK1 and PKM1 are allosterically activated by AMP and may up-regulate glycolysis to produce more ATP. | 154,155 |
|  | Glycogen metabolism | To meet the energy demands of neurons, metabolism of glycogen to glucose in astrocytes may be enhanced. | 156,157 |
| Post complex V | Creatine de-phosphorylation | Free ATP is produced when creatine is de-phosphorylated. This is limited by availability of phosphorylated creatine. | 149 |
|  | Lipid transfer | Excess fatty acids generated under high energy demands would be transported to astrocytes to limit further ROS. | 158 |

#### 3.5 Metabolic comparison between terminally differentiated cell types

**Table 5:** Comparison of metabolism between neurons, astrocytes and muscle cells. All these cells are terminally differentiated.

| *Process* | *Neurons* | *Astrocytes* | *Type 1 muscle* | *Type 2A muscle* |
| --- | --- | --- | --- | --- |
| Nutrient source | BBB, astrocytes | BBB, capillary | capillary | capillary |
| Aerobic | yes | yes | yes | yes |
| Anaerobic | no | yes | no | yes |
| Oxidative phosphorylation | yes | low | yes | yes |
| Primary fuels | lactate, glucose | glucose | glucose | glucose |
| Fatty acid fuels | yes | long-chain | yes | yes |
| Amino acid fuels | yes | yes | yes | yes |
| Ketone bodies | yes | yes | yes | yes |
| Creatine | yes | yes | low | yes |
| Carbohydrate reserves | none | glycogen, lipids | triglycerides | glycogen |
| O<sub>2</sub> homeostasis | neuroglobin | neuroglobin | myoglobin (high) | myoglobin (low) |
| Primary glucose intake | GLUT3 | GLUT1 | GLUT4 | GLUT4 |
| Extracellular glucose conc. | 2-3mM | 2-3mM | 6-7mM | 6-7mM |
| Insulin-glucose uptake | largely insensitive | largely insensitive | yes | yes |
| Metabolic end products | ATP | lactate, ATP, glutamine | ATP | ATP, lactate |
| Primary ATP consumer | ion pumps | ion pumps, glutamine synthetase | myosin ATPase | myosin ATPase |
| Multi-nucleated | no | no | yes | yes |

## 3.6 Acronyms and Abbreviations

*α*KG: Alpha ketoglutarate
*α*Syn: Alpha synuclein
CoQH_2_/CoQ: Coenzyme Q (reduced/oxidized form)
FADH_2_: Flavin adenine dinucleotide (reduced)
IP_3_: Inositol 1,4,5-trisphosphate
PIP_2_: Phosphatidylinositol 4,5-bisphosphate
6PG: 6-P-gluconate
Act.CoA: Acetyl coenzyme A
ADP: Adenosine diphosphate
ADPR: Adenosine diphosphate ribose
AIS: Axon initial segment
ALT: Alanine transaminase
AMP: Adenosine monophosphate
ANT: Adenine nucleotide translocator
AOX: Alternative oxidase
APC: Anaphase-promoting complex
AST: Aspartate transaminase
ATP: Adenosine triphosphate
BBB: Blood brain barrier
cAMP: Cyclic adenosine monophosphate
Cit: Citric acid
CMA: Co-chaperone mediated autophagy
CV: Coeffecient of variance
D_*C*_, D_*E*_, D_*V*_: Dopamine cytosolic, extracellular, vesicle
DA-Q: Dopamine-derived quinones
DAG: Diacyl glycerol
DAT: Dopamine transporter
DHAP: Dihydroxyacetone phosphate
DHODH: Dihydroorotate dehydrogenase
DOPAC: 3,4-dihydroxyphenylacetic acid
DOPAL: 3,4-Dihydroxyphenylacetaldehyde
ETC: Electron transport chain
ETF: Electron-transferring flavoprotein
F1,6BP: Fructose 1,6-bisphosphate
F2,6BP: Fructose 2,6-bisphosphate
F6P: Fructose 6-phosphate
FETROS: Forward electron transport ROS
G3P: Glycerol 3-phosphate
G6P: Glucose 6-phosphate
GA3P: Glyceraldehyde 3-phosphate
GBA: Glucocerebrosidase
GDH: Glutamate dehydrogenase
GEF: Guanine nucleotide exchange factors
GPCR: G protein-coupled receptors
GPDH: Glycerol-3-phosphate dehydrogenase
Gpe: external globus pallidus
Gpi: internal globus pallidus
GS: Glutamine synthetase
GSH: Glutathione (reduced)
GSSH: Glutathione disulphide (oxidized)
GTP: Guanosine triphosphate
HEX: Hexokinase
HK1: Hexokinase-1 enzyme
HSC70: Heat shock cognate 71 kDa protein
IDH: Isocitrate dehydrogenase
ISI: Interspike interval
LDH1/5: Lactate dehydrogenase type 1 or 5
LRRK2: Leucine-rich repeat kinase 2
MCU: Mitochondrial calcium uniporter
ME: Malic enzyme
MIM: Mitochondrial inner membrane
miniSOG: small and efficient singlet oxygen generator
MOM: Mitochondrial outer membrane
MPP+: 1-methyl-4-phenylpyridinium
MS: Metabolic signal
NADH/NAD+: Nicotinamide adenine dinucleotide (reduced/oxidized)
NADPH/NADP+: Nicotinamide adenine dinucleotide phosphate (reduced/oxidized)
NCX: Sodium calcium exchanger
Oxa: Oxaloacetate
PC: Pyruvate carboxylase
PDH: Pyruvate dehydrogenase
PDK4: Pyruvate dehydrogenase kinase 4
PDP: Pyruvate dehydrogenase phosphatase
PEP: Phosphoenolpyruvic acid
PEPCK: Phosphoenolpyruvate carboxykinase
PFK: Phosphofructokinase
PFKFB3: 6phosphofructo2kinase/fructose2,6biphosphatase3
PINK1: PTEN-induced kinase 1
PKA: Protein kinase A
PKC: Protein kinase C
PKM1: Pyruvate kinase M1
PLC: Phospholipase C
PPP: Pentose phosphate pathway
R5P: Ribose 5-phosphate
RETROS: Reverse electron transport ROS
ROS: Reactive oxygen species
SCNA: Alpha synuclein
SNcDA: Substantia nigra pars compacta dopaminergic neuron
SNr: Substantia nigra pars reticulata
SOD2: Superoxide dismutase
STN: Subthalamic nucleus
TCA: Tricarboxylic acid
TXN: Thioredoxin
UCP: Uncoupler protein
USP: Ubiquitin-specific protease
V-ATPase: Vacuolar-type ATPase
VMAT2: Vesicular monoamine transporter 2

## 4 Supplementary Figures

### 4.1 Physiological factors that affect ROS relief

**Figure S1:**
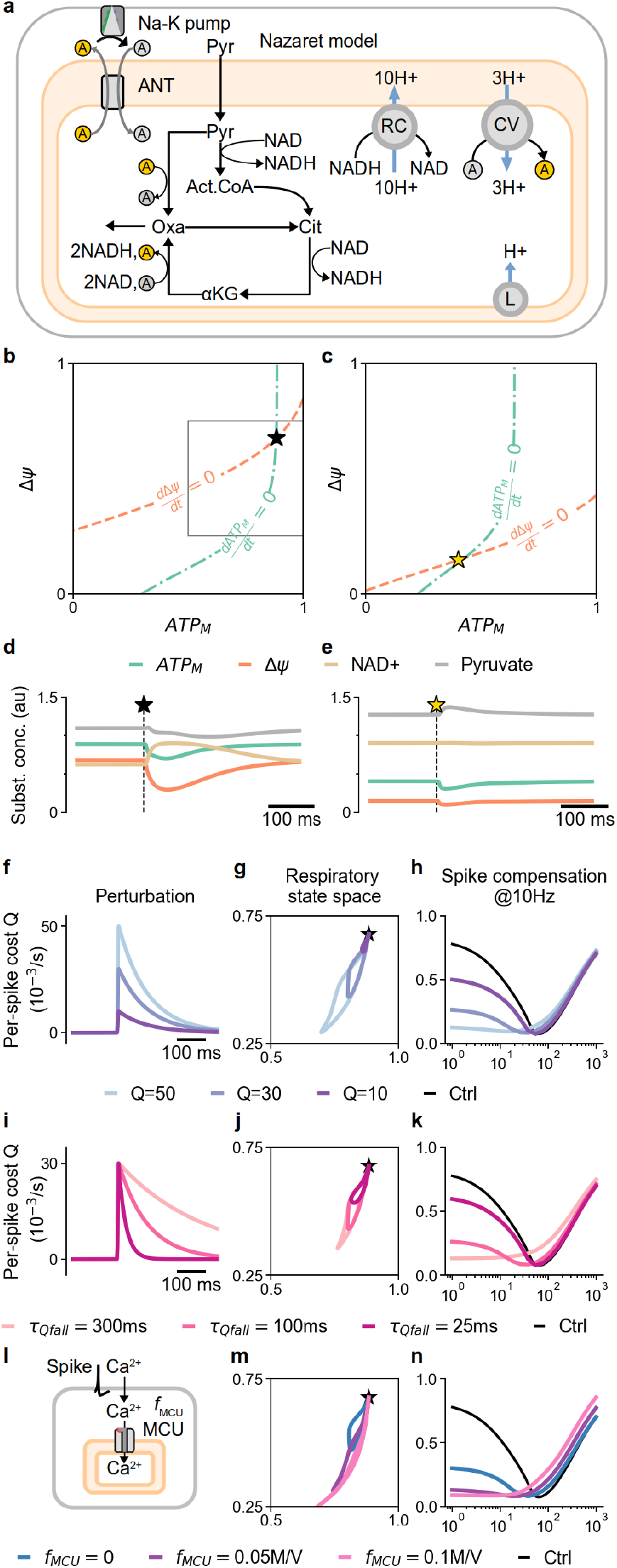
**a)** A schematic of a simplified TCA/ETC model for mitochondrial ATP production, i.e., the Nazareth model^54^. Here the rate at which ATP_*M*_, and ADP_*C*_ are exchanged to ADP_*M*_ and ATP_*C*_ at the adenine nucleotide translocator (ANT), is given by the rate constant *k*_*ANT*_ and determines the steady state values of all substrates (in CAPS). **b**,**c)** *k*_*ANT*_ at 30 /ks (black star) and 150 /ks (gold star) corresponds to the fixed points of the nullclines of ATP_*M*_ (green) and ΔΨ (orange). **d**,**e)** Changes in the concentrations of all mitochondrial substrates (ATP_*M*_ (green), ΔΨ (orange), NAD+ - oxidised form of NADH (tan) and Pyruvate (gray, Pyr)) in response to a single spike of cost Q (30 /ks in *k*_*ANT*_).**f)** Examples for perturbations with various values of per-spike cost Q. **g)** Excursions in the respiratory space (inset from b) following perturbations shown in f, starting from the black star. **h)** ROS level compensation due to metabolic spiking at 10 Hz with various Q. **i)** Examples for perturbations with various recovery time constants. that change the respiratory steady state excursion **j)** and the amount of ROS relief due to metabolic spiking at 10 Hz **k. l)** Schematic of Ca2+ entering the mitochondria following a spike. m) Resulting ΔΨ depletion, modelled as an increasing the H+ leak (see a, label L). **m)** ROS relief as a function of Ca2+ entry. **n)** ROS level compensation due to metabolic spiking at 10 Hz for various levels of Ca2+ entry.

### 4.2 Time course of ROS relief

**Figure S2:**
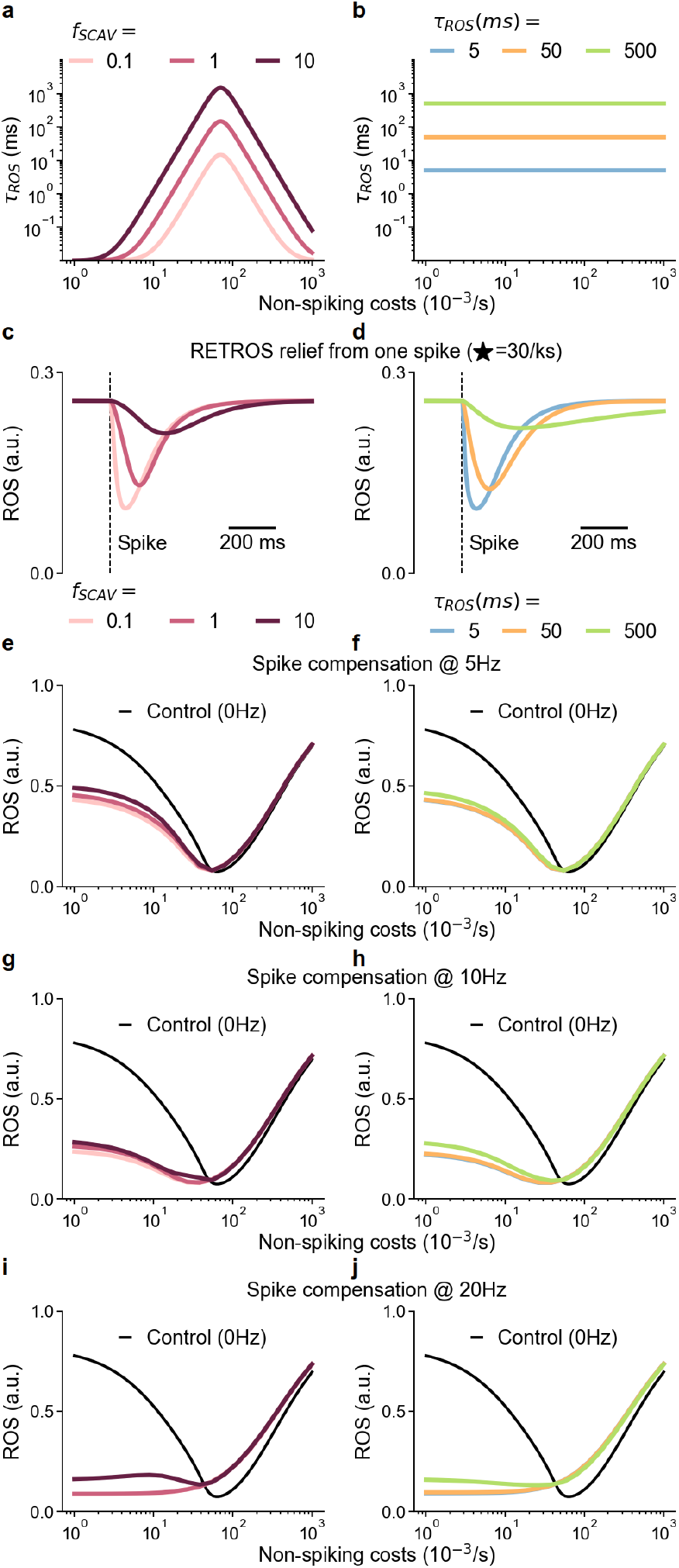
The time constant of the ROS accumulation in a neuron depends on its ROS scavenging capacity and may vary depending on the neuron type. Due to lack of experimental data, we explored a large range of time scales *τ*_*ROS*_ for ROS scavenging effectiveness, either **a)** as a non-spiking costs dependent function with maximum effectiveness near ROS minimum (pink, red and maroon) or as **b)** a constant value (blue, orange and green). The corresponding changes in RETROS relief per spike are show in **c** and **d**. A single spike occurs at the dotted vertical line. The overall ROS levels change due to spiking at 5 Hz **(e**,**f)**, 10 Hz **(g**,**h)** and 20 Hz **(i**,**j)**. These results suggest that for a wide range of time scales of ROS accumulation, spiking can effect ROS levels in neurons. Importantly, spikes can provide ROS relief as seen by the collapse of the RETROS levels in plots **e-j**.

### 4.3 Summary plots for FETROS and RETROS responses in the accounting model

**Figure S3:**
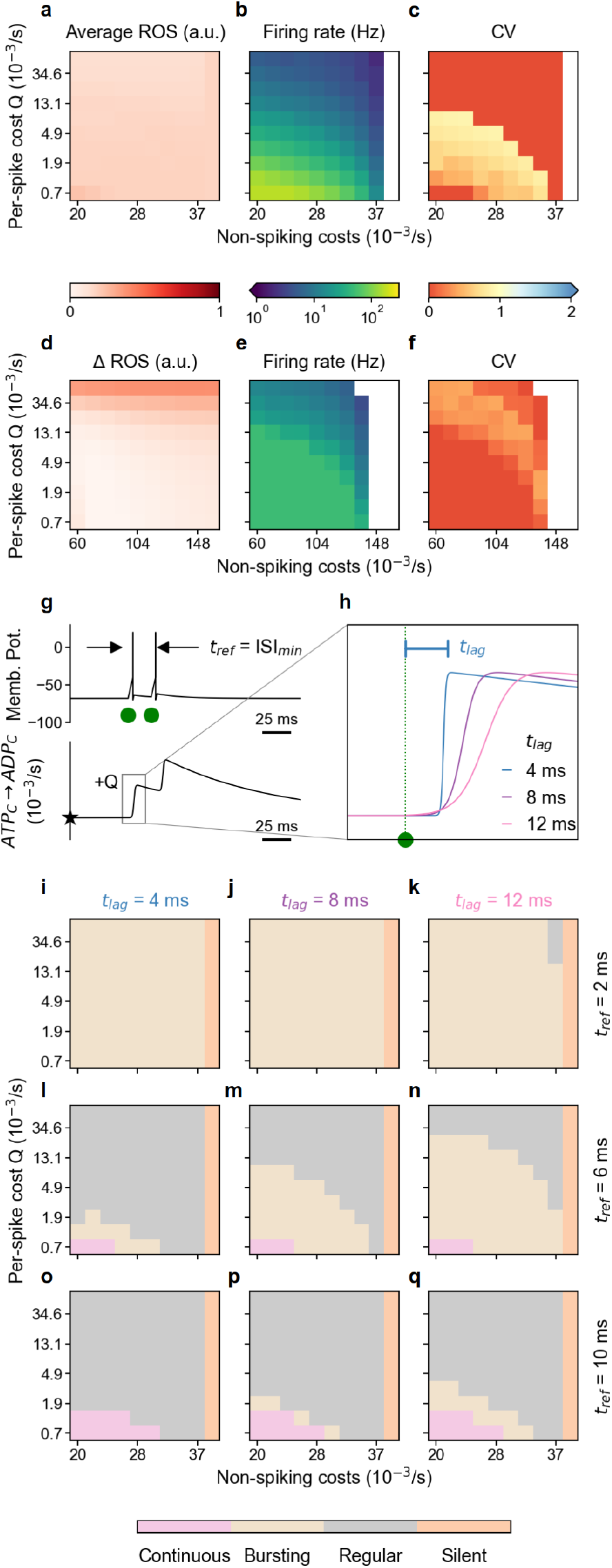
**a-c)** RETROS response: **a)** Low average ROS levels for metabolic spiking at different levels of baseline ATP consumption and cost-per-spike Q as described in Fig. 2e. **b)** Metabolic firing rate maintaining low ROS levels and **c)** coefficient of variance of the inter-spike intervals (CV ISI) for each parameter pair in a). **d-f)** FETROS response when additional spikes are induced (at 44 Hz and with a CV = 1). **d)** Difference in ROS levels between models with and without a FETROS response. **e)** Resulting firing rates, and **f)** CV ISI for each parameter pair in **d** as described in Fig. 2f. **g)** Schematic of two spikes (top) and the resulting transient increases in ATP expenditure (bottom). Green dots denote spike threshold crossing. The minimum interval between crossings is denoted as the refractory period (refrac). **h)** Detail of the rise time of ATP consumption after spike initiation (green dotted line) according to per-spike cost Q. **i-q** Metabolic spike patterns as a function of baseline non-spiking costs (x-axes), per-spike costs Q (y axes), refractory period (2 ms for **(i**,**j**,**k)**, 6 ms for **(l**,**m**,**n)**, 10 ms for **(o**,**p**,**q)** and time-to-peak of the metabolic response (4 ms for **(i**,**l**,**o)**, 8 ms for **(j**,**m**,**p)**, 12 ms for **(k**,**n**,**q)**).

### 4.4 Summary plots for metabolic recurrent network model

**Figure S4:**
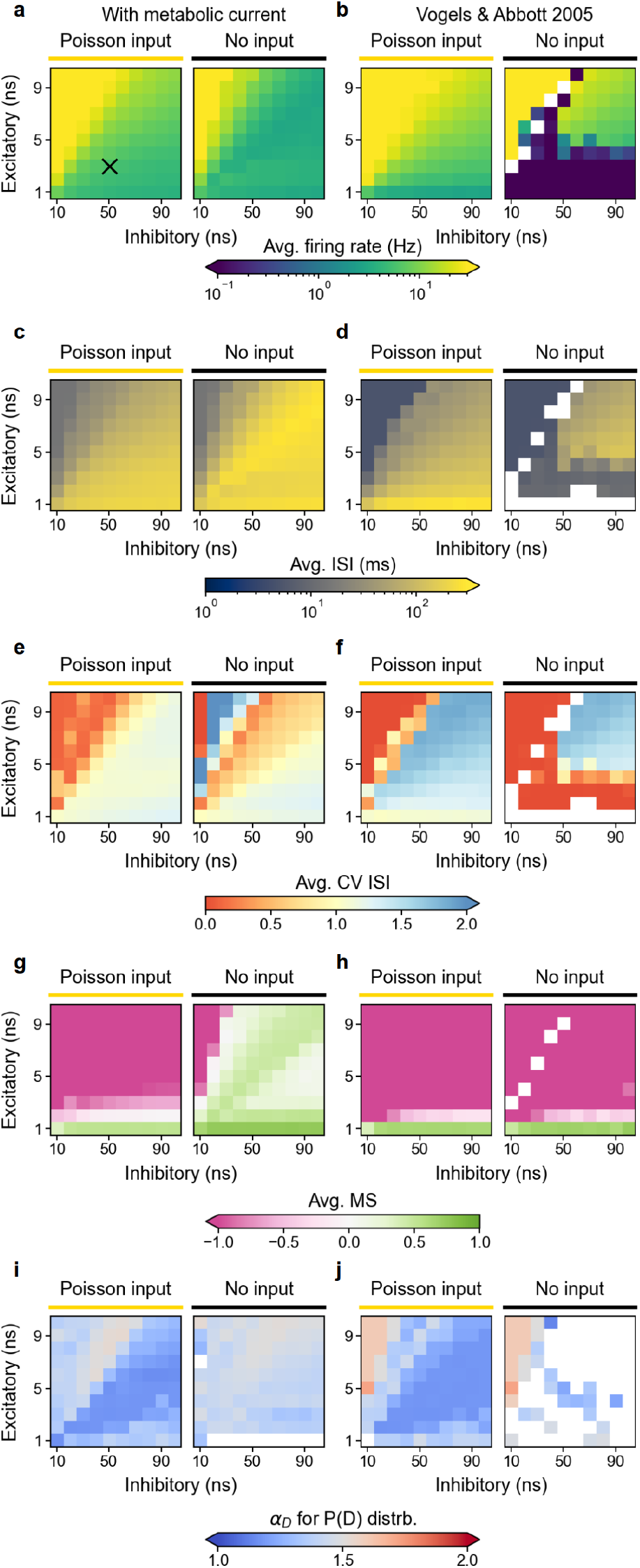
**a, c, e, g, i)** Recurrent network model with metabolic current compared to the model without **b, d, f, h, j)**. All the values shown here correspond to the averages as indicated by the dotted lines in Fig.3 l,m,n,o, and slope of the line in Fig.3p: **a, b)** Average firing rates over 5 s simulated time of the network models as a function of inhibitory (x-axes) and excitatory (y-axes) conductance for the cases driven by an external Poisson input of 3 Hz (left), and without any external input (right) The black x in the left most plot indicates the parameter pair used for simulations in Fig. 5. **c, d)** Average inter-spike intervals, **e, f)** average coefficient of variance of the inter-spike intervals (CV ISI) and **g, h)** the average metabolic signal at the time of spiking. **i, j)** The slope of the distribution of avalanche durations (1 ms time bins). White patches in these colormaps indicate insufficient data.

### 4.5 Known co-factors leading to PD and their placement in our framework

**Figure S5:**
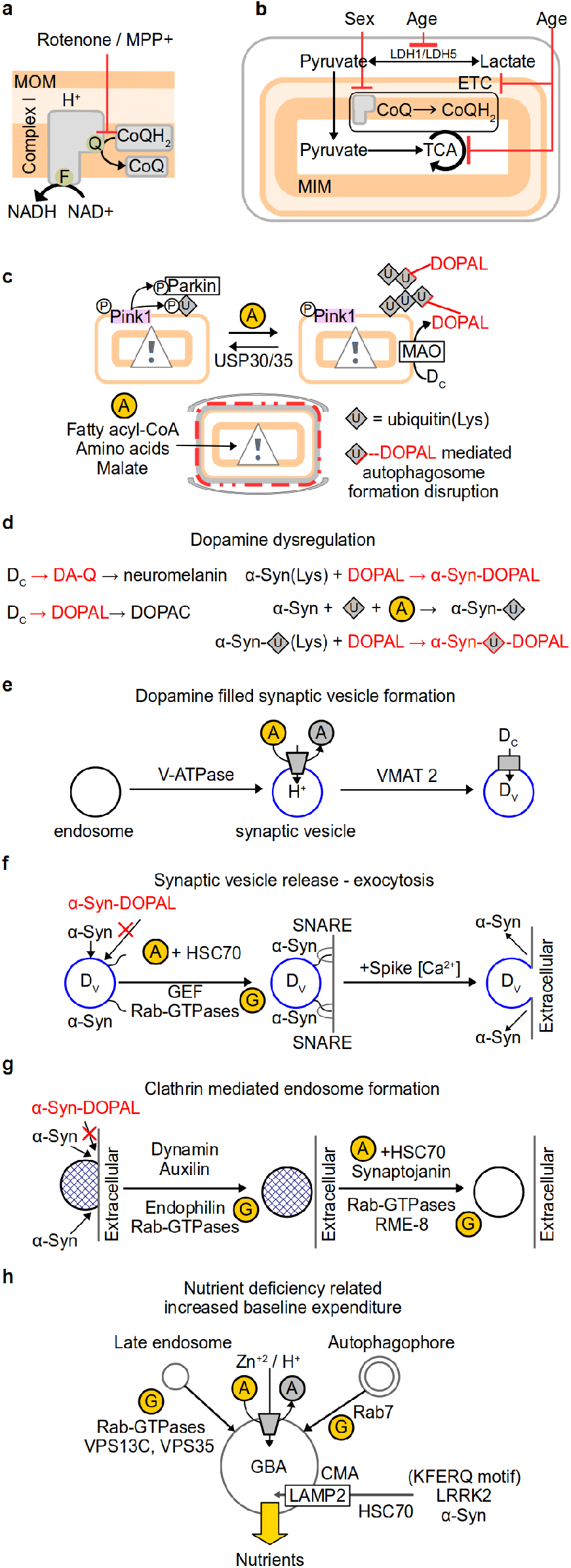
We assigned many well-identified co-factors leading to PD due to lack of dopamine in the striatum to the three broad categories of homeostatic defects of metabolic spike initiation in SNcDA (Fig. 6). Red corresponds to defects in the RETROS signalling cascade, blue corresponds to increases in per-spike cost, and gold (including ATP, GTP circles) corresponds to an overall increase in non-spiking related expenses. **a)** Chronic exposure to rotenone or MPP+ may cause competitive binding at the ubiquinone binding site of complex I, blocking RETROS release site. **b)** Age and sex related co-factors may cause inefficient mitochondrial ATP production. **c)** Defects in tagging, trafficking and removal of mitochondria marked for repair or mitophagy may cause resource competition with their healthy counterparts. **d)** Lack of extracellular dopamine (D_*E*_) release may drive cytosolic dopamine (D_*C*_) dysregulation to produce DOPAL and quinones, and further deleterious side-effects by their interaction with lysine (Lys) residues. **e)** In SNcDA neurons, dopamine is also released from the somatodendritic compartments. Any faults in filling synaptic vesicles with dopamine (D_*V*_), or **f)** failures in exocytosis of vesicle pools may lead to increased per-spike expenditure. Deficiencies in energy dependent **g)** clathrin mediated formation of endosomes, which are essential for synaptic vesicles’ and lysosomal function, may increase non-spiking costs. Similarly, **h)** defects in lysosomal formation or in delivering cytosolic components marked for degradation to the lysosome may result in aggregates in the cytosol and subsequent lack of nutrients - in effect also increasing the baseline non-spiking expenditure.

### 4.6 Overview of neuronal glucose metabolism

**Figure S6:**
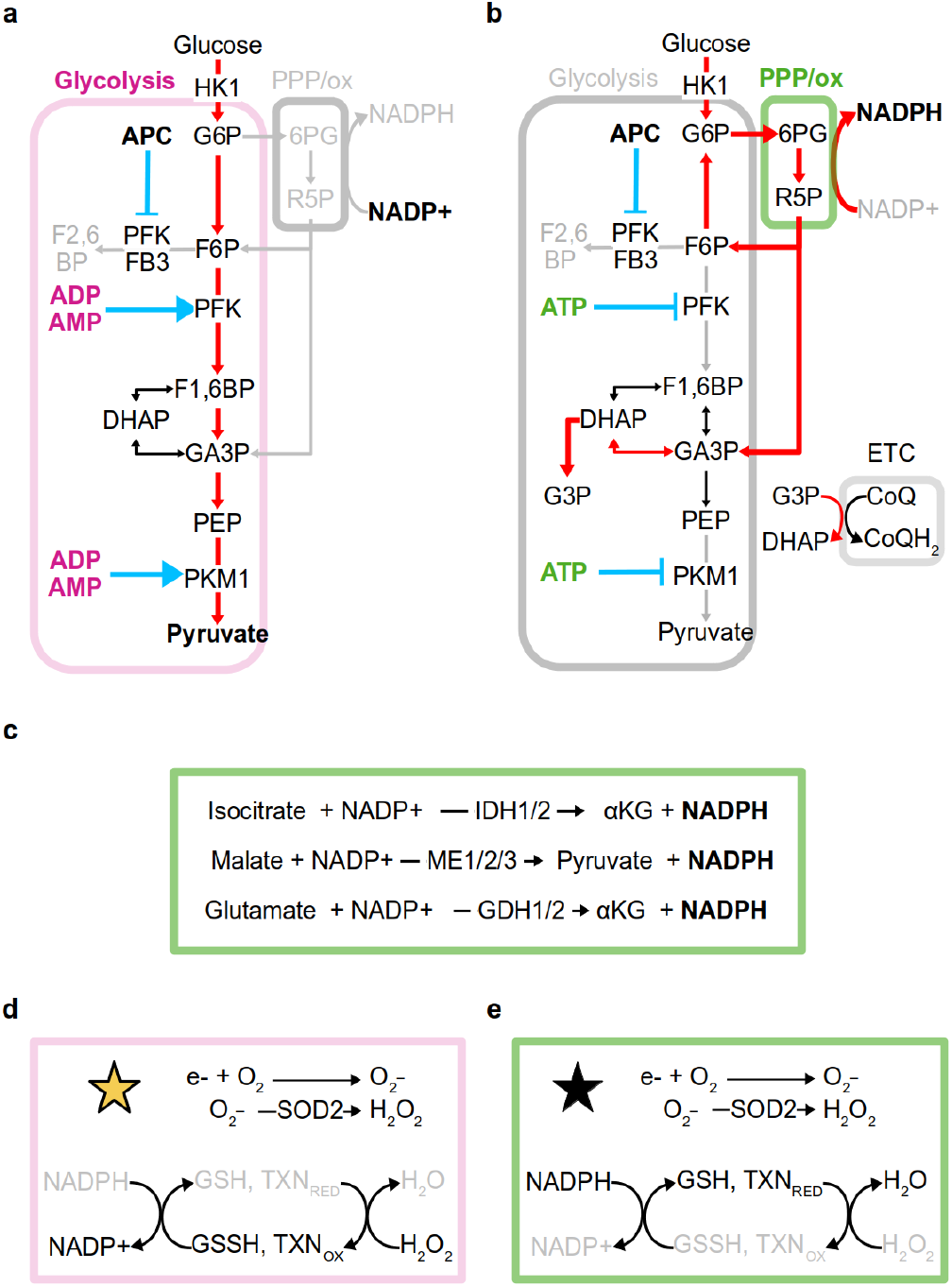
**a**,**b)** Neuronal glucose metabolism in RETROS and FETROS conditions, respectively. The preferred pathways for glucose breakdown are shown in red. Neuron-specific regulation is indicated in blue. Unlike in other cell types, the expression of APC inhibits PFKFB3, lowering available F2,6BP. Consequently neuronal PFK and PKM1 isoform are primarily regulated by ATP/ADP/AMP. **a)** Under FETROS, ADP/AMP up-regulate glycolysis (pink box) and glucose is diverted away from the PPP/ox (gray box), limiting NADPH production (gray). **b)** Under RETROS conditions, glucose is routed through the oxidative phase of the pentose phosphate pathway (PPP/ox, green box) instead of glycolysis. Consequently NADPH production is up-regulated (bold), and glycolysis is down-regulated (indicated in gray) by high ATP (bold). **c)** Other reactions that can produce NADPH may also be up-regulated under RETROS. **d, e)** Mitochondrial ROS is scavenged in the cytosol by redox couples GSH/GSSG, thioredoxin (TXN), etc., which are replenished by NADPH. Low and high concentrations are indicated in gray and black respectively. **d)** Under FETROS (gold star), NADPH supply is limited and consequently the redox pools are mostly oxidized. **e)** Under RETROS conditions (black star) the scavenger pools operate at their maximum capacity and are mostly reduced.

### 4.7 Overview of neuronal metabolic flexibility

**Figure S7:**
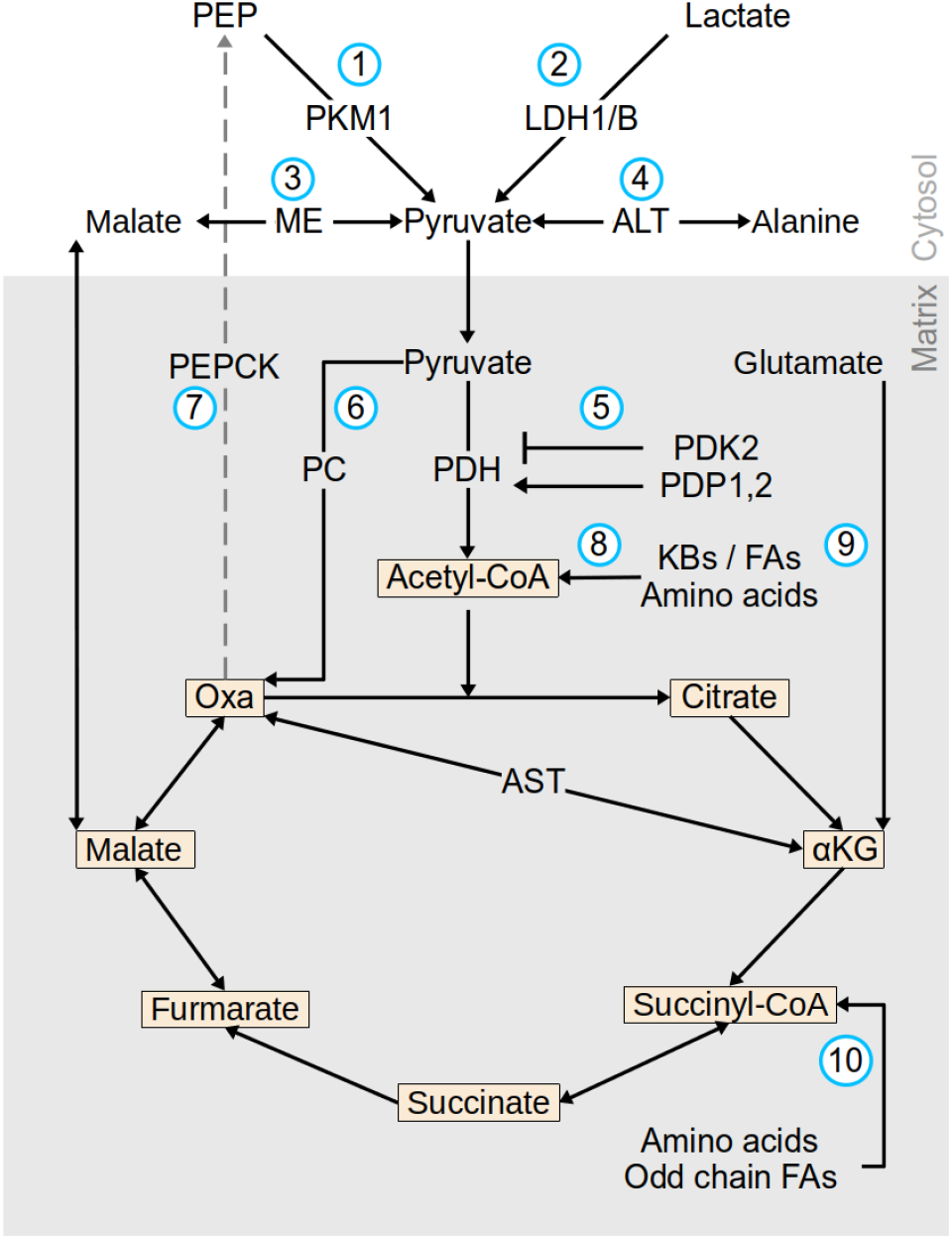
Metabolic pathways in neurons may operate flexibly to percolate metabolic fuels perpetually into the mitochondria, thus maintaining a high ΔΨ. Neuron-specific pathways are indicated in blue circled numbers and citric acid cycle (TCA) substrates are shown in peach colored boxes. Cytosolic pyruvate in neurons can be produced from glycolysis, where phosphoenolpyruvate (PEP) is converted to pyruvate. In neurons, this step is regulated by PKM1 isoform (1) which lacks allosteric regulation by F1,6BP and alanine. Pyruvate may also be produced from lactate (2), from malate (3) and from some amino acids such as alanine (4). Mitochondrial pyruvate enters the TCA as acetyl-CoA by the pyruvate dehydrogenase (PDH). In neurons, PDH perhaps largely remains in its active state due to low expression of its inhibitory regulator (5) pyruvate dehydrogenase kinases (PDK1,3,4) and high expression of its activator pyruvate dehydrogenase phosphatase (PDP1,2). Pyruvate may also enter the TCA indirectly as (6) oxaloacetate (Oxa). Oxa (and glutamate) can be converted to aspartate (and *α*KG) instead of PEP due to limited phosphoenolpyruvate carboxykinase (PEPCK) expression (7, gray dashed line). Acetyl-CoA itself can also be augmented by (8) ketone bodies (KBs), fatty acids (FAs) and amino acids. Additionally glutamate (9) - the most abundant free amino acid in the brain, can also enter TCA via *α*KG. Finally, odd chain FAs and some amino acids (10) can also produce TCA intermediate succinyl-CoA.

## Footnotes

† with instantaneous ATP-ADP exchange with the cytosol, a weak assumption under FET, when other sources of ATP like glycolysis or creatine phosphorylation may also be utilised

†† A weak assumption under FET where Ca_2_^+^ buffering and up-regulation of TCA+ETC in neurons is anticipated^<sup>55</sup>^

§ but note that *f*_*Q*_ increased when we modelled demyelination or mitochondrial migration away from the spike initiation micro domain

* Notably, acute exposure to rotenone such as during patch clamp experiments has a different response.^60,61^

## Notes

### Competing Interest Statement

The authors have declared no competing interest.

